# Tumor-Educated-Platelets interact with Breast Cancer-Stem-Cells via P-selectin- PSGL1 and ensure stemness and metastasis through WNT-β-Catenin-VEGF-VEGFR2 intra-cellular signaling: Therapeutic modulation by aspirin

**DOI:** 10.1101/2025.06.04.657784

**Authors:** Aishwarya Guha, Jasmine Sultana, Avishek Bhuniya, Mohona Chakravarti, Saurav Bera, Anirban Sarkar, Sukanya Dhar, Pritha Roy Choudhury, Juhina Das, Nilanjan Ganguly, Ipsita Guha, Tapasi Das, Neyaz Alam, Indranil Ghosh, Srabanti Hajra, Rathindranath Baral, Anamika Bose, Saptak Banerjee

## Abstract

**Background:** Protagonistic role of platelets promote capillary infiltration of tumors for distant metastasis along with immunosurveillance. Despite existing reports highlighting role of platelets in tumorigenesis, its impact on breast cancer stem cells (BCSCs) remain underexplored. Our first ever report on murine and human system, accentuate that, tumor educated platelets (TEPs) of luminal A and TNBC subtypes are distinct from healthy counterparts, collaborating with BCSCs to generate sub-variants that elevate tumor aggressiveness.

**Methods:** Impact of TEPs on BCSCs was evaluated from primary breast tumor and blood samples of luminal A/TNBC patients along with EC/4T1 murine breast tumor models and MCF-7/MDA-MB-231 cell lines. For downstream assays, TEPs were co-cultured with breast tumor samples or cell lines, followed by magnetic sorting of lin^-^CD44^+^CD24^-^ BCSCs. TEP induced alterations of BCSCs were evaluated from 3D tumorsphere, colony formation, transwell migration, scratch-wound healing, matrigel invasion, *in-vitro* tube formation assays. Fluorescence-confocal microscopy, RT-PCR, flow-cytometry, western-blotting was utilized to decipher the role of genes and protein involved in stemness, metastasis along with the transcription factors in the downstream signaling cascade, followed by verifications by RNAi.

**Results:** TEPs have elevated expression of P-selectin and interacts with BCSCs via P-selectin and PSGL1 on BCSCs surface. Treatment with aspirin had restorative impact on P-selectin level, converting TEPs from active to resting platelet (RP) state. Under TEPs influence, BCSCs were tumorigenic, clonogenic, multidrug resistant, invasive with numerous invadopodia and remained skewed towards mesenchymal phenotype. Administration of RP reduced TEP associated BCSC virulence both *in-vivo* and *in-vitro.* P-selectin-PSGL1 interaction results in binding of WNT to FRIZZLED followed by stabilization and nuclear translocation of β-Catenin. Nuclear β-Catenin promotes stemness-EMT (Epithelial to mesenchymal transition)- metastasis, along with stimulation of autocrine VEGF-VEGFR2 cascade. Inhibition of WNT and VEGFR2 by RNAi confirmed the critical role of this axis in regulating TEPs influence on BCSCs.

**Conclusion:** These insights into TEPs-BCSC interplay, acknowledges TEPs, as-well-as unveils novel receptor-ligand signaling cascade between TEPs and BCSCs, that could be a beneficial therapeutic strategy to target cancer metastasis.

**Graphical Abstract:** 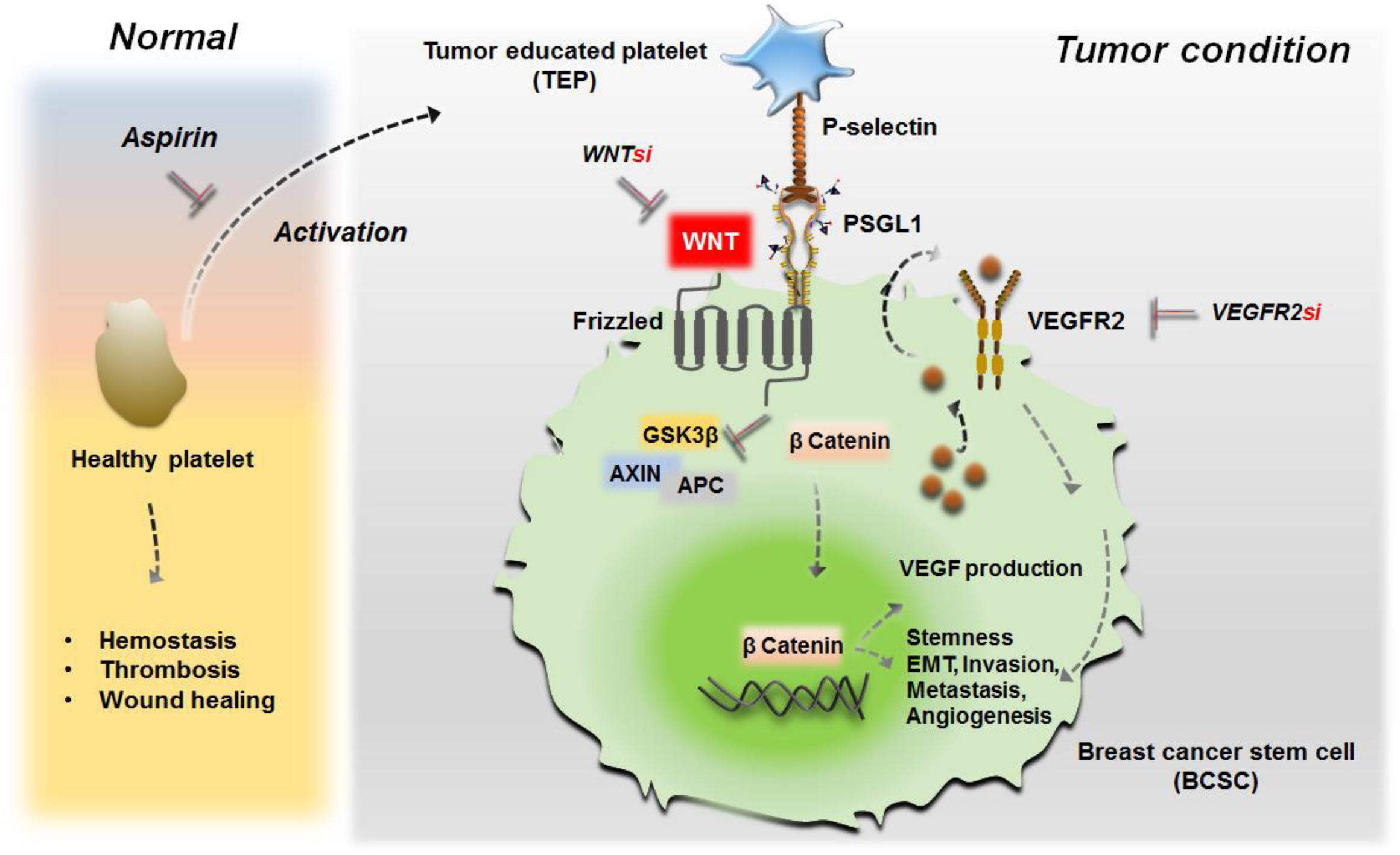

Cellular interaction between TEPs P-selectin and BCSCs PSGL1 to regulate stemness, EMT, metastasis and angiogenesis via WNT-β-Catenin-VEGF-VEGFR2 cascade: Modulation by pharmacological inhibitor and RNAi.

## Introduction

Based upon the expression of hormone receptors, breast cancer (BC), the most common type of cancer among women globally, can be categorized into four main subtypes (1–3) luminal A, luminal B, Her2^+^ and triple negative (TNBC), making TNBC most aggressive subsequently in the order (1–3). The rapid spread of BC to distant organs or lymph nodes, makes therapy difficult, contributing to the disease’s aggressiveness and poor prognosis (4).

A potential explanation for this aggressiveness is the existence of a rare population of quiescent, infrequently dividing cells called cancer stem cells (CSCs) (5–6). The CSC protective niche within the tumor microenvironment (TME) along with high expression of drug efflux proteins makes them ‘immortal’ and challenging to target (7), shielding them from the effects of therapeutic drugs (7). Additionally, they trigger the extracellular matrix’s (ECM) breakdown, which makes invasion and metastasis easier (7–8).

Metastasis accounts for majority of cancer-related mortalities worldwide (9). Nevertheless, the ability to successfully develop secondary tumor is limited to only a small population of cells from the main tumor bulk (10). During their voyage through the circulation, these cells are capable of evading the ‘immunosurveillance process’ by the help of tiny anucleated platelets present in the blood stream (11–13). Being the second most abundant cell type of the peripheral blood, platelets have emerged as central players of EMT and metastasis (14–15).

Reciprocal interaction between the tumor cells and platelets cause activation of the latter which are termed as ‘tumor educated platelets’ or TEPs (16–19). Subsequent studies revealed that TEPs differ significantly from normal counterpart, producing numerous filopodial process that facilitates sequestration and protection of the tumor cells. Additionally, they are a repository of various biologically active molecules that influence tumor progression by modulating EMT, metastasis and angiogenesis (20).

Several reports have mentioned about the role of TEPs in advancing EMT and metastasis in solid tumors including BC through cell surface receptor-ligand interactions (21–23). However, the status and function of TEPs within the TME has remained under-explored. Our study has found that TEPs are a pivotal component of the breast TME and interact with breast cancer stem cells (BCSCs) through physical contact. P-selectin on the surface of TEPs binds with PSGL1(P-Selectin Glycoprotein Ligand 1) on the surface of BCSCs, facilitating the clustering of TEPs around these cells.

This coaction indicated towards their possible alliance in promoting tumor progression. Through numerous *in-vitro* and *in-vivo* experiments, we have found that TEPs directly interact with BCSCs generating highly aggressive, clonogenic, therapy resistant, migratory and invasive BCSC sub-variants that encouraged pulmonary metastasis of BC *in-vivo* via WNT-β-Catenin-VEGF-VEGFR2 axis in the downstream. This implies the importance of therapeutically targeting TEPs to control BC progression. Pharmacologically inhibiting TEPs with aspirin not only reduced its activation but also its BCSC proliferating effect. Additionally, by neutralizing WNT-β-Catenin-VEGF-VEGFR2 cascade the aggressiveness of the disease could be compensated.

## Methods

### Reagents and antibodies

MEM, RPMI-1640, DMEM high-glucose, DMEM: F12K (1:1), foetal bovine serum (FBS) were purchased from Hi-Media (Mumbai, India). B27^TM^ supplement (50X) was purchased from Gibco (Waltham, USA). Recombinant human epidermal growth factor (rEGF) and recombinant human basic fibroblast growth factor (rbFGF) were purchased from Merck-Milipore (Damstadt, Germany), Aspirin, Fluorshield ^TM^ DAPI was procured from Sigma Aldrich (Missouri, USA). Detailed list of all antibodies utilized is provided in the Supplementary Table T-1.

### Human solid tumors

Pre-treatment and post-operative breast tumor samples (luminal A n=20, TNBC n=20) and with proven histopathological normalcy were collected from Chittaranjan National Cancer Institute (CNCI), Kolkata, India following approval from patients and Institutional Ethical Committee (Approval no: CNCI-IEC-SB-2020-30). TNM staging status was noted during collection.

### Human blood

Blood (5ml) was procured from pre-treatment and post-operative breast carcinoma patients (n=40) by venepuncture method from Chittaranjan National Cancer Institute (CNCI), Kolkata, India following approval from patients and Institutional Ethical Committee (Approval no: CNCI-IEC-SB-2020-30). 5ml blood was also collected from healthy donors who were not under any medications and did not consume any platelet lowering drugs for atleast 1month were used as controls. Patient details are provided in Supplementary Table T-3.

### Inclusion and exclusion criteria for human tumor and blood samples

Pre-treatment and post-operative breast tumor and blood samples were collected from patients above 18 years and below 70 years age. Patients with any prior history of co-morbidity were excluded from the study. Blood was not collected from any patients who consumed platelet lowering drugs (aspirin) for atleast 1 month.

### Isolation of tumor educated platelets (TEPs)

TEPs were isolated from whole blood by double centrifugation method. Briefly 5ml blood was collected in ACD anticoagulant and initially centrifuged at 260g x 20 minutes. This separates the straw-coloured platelet rich plasma (PRP) from the underlying RBC zone. The PRP was separated to two parts. One part was untreated and to the other part 100µm/ml aspirin was added for 30 minutes to restore the resting condition of platelets (resting platelets-RP). The PRP was further centrifuged at 800g x 20 minutes without brakes to separate platelet pellet from platelet poor plasma (PPP). The pellet was then washed with platelet wash buffer thrice and dissolved in Tyrode buffer and was ready for use.

### SEM Microscopy

Ultrastructure of platelets were visualized by SEM microscopy. Approximately 20μL of PRP was taken on coverslip and placed on moist filter paper and incubated at 37°C for 5 minutes. This allowed the PRP to dry in a humid environment. Post incubation, the samples were washed with 1X PBS to remove excess plasma. Samples were fixed in 2.5% glutaraldehyde for 30 minutes followed by triplicate washing with 1 X PBS for 5 minutes each. Further fixation was done in 1% osmium tetroxide for 1 hour. The samples were thoroughly washed and dehydrated in ascending grades of ethanol starting with 30%, 50%, 70%, 90% and finally 100% with atleast 15 minutes incubation at each step. Samples were then transferred to carbon tapes on aluminium stubs, gold coated and visualized using Zeiss Gemini Scanning Electron Microscope.

### Cell lines and culture

MCF-7 (RRID: CVCL_0031), MDA-MB-231 (RRID: CVCL_0062), 4T1 (RRID: CVCL_0125) cell lines were obtained from National Centre for Cell Sciences (NCCS), Pune, India. Cells were cultured in complete media supplemented with 10% (v/v) heat inactivated FBS, 2mM L-glutamine, 100 U/ml penicillin and 100µg/ml streptomycin in incubator at 37^°^C and 5% CO^2^. Cell lines were monitored regularly for any changes in morphology. Mycoplasma contamination if any was removed by EZkill^TM^ Mycoplasma Elimination kit procured from Himedia (Mumbai, India). Cells were maintained for 10-12 passages and all experiments were performed within 6 months of purchase.

### Co-culture

Single cell suspensions of luminal A and TNBC tumor samples and MCF-7, MDA-MB-231, 4T1 and EC were cultured with TEPs and RP in 1:100 (BCSC: platelet) ratio for 24 hours. After the incubation period, the culture media with platelet suspension was removed and the cells were collected by trypsinization and centrifuged for further analysis.

### Magnetic cell sorting

From the co-culture setup BCSCs were sorted by using magnetically labelled CD44^+^/CD24^-^/Lineage^-^-cocktail antibodies and cell purification was carried out according to manufacturer’s protocol (MicroBead kit, Miltenyi Biotech, Germany). Purity of cells was checked by flow cytometry.

### CSC enrichment culture and tumorsphere assay

CSC enrichment media was prepared by supplementing serum free DMEM: F12K (1:1) media with 1% B27^TM^ supplement (50X). To it heparin (40ng/mL), human-rEGF (20ng/mL), human-rbFGF (20ng/mL) were added freshly. CSCs (1x10^4^) were cultured in this media and plated on ultra-low adherent plate (Corning, New York, USA). Cells were incubated in 5% CO_2_ humified atmosphere at 37^°^C for 7 days and fresh media was supplemented to the culture after every three days. Tumorspheres were micrographed and their count from 5 random fields was documented. Their area was calculated using ImageJ software. Tumorspheres so formed were dissociated with trypsin and were centrifuged to produce single cell suspensions which were further analysed.

### Mice and tumors

Wild-type female BALB/c and Swiss albino mice (*Mus musculus*) (age: 4-6 weeks; body weight: 18-22g average) obtained from Institutional Animal Care and Maintenance Department were included in the experiments. Mice that showed signs of illness or stress during acclimatization were proposed to be excluded from the study. Since, the animals did not appear to show any such symptoms, all of them were included in the investigation. Animals were maintained in pathogen free environment and fed with autoclaved food (Epic Laboratory, West Bengal Government, Kalyani, India) and water *ad libitum.* All experiments were performed after approval from Institutional Animal Care and Ethics Committee (Approval No: IAEC-1774/SBn-4/2021/9). 4T1 cells were maintained *in-vitro* and EC cells (Ehrlich Carcinoma) were maintained in swiss albino mice as intraperitoneal passage. To minimize biasness and controlling of biological variability, mice were allocated in the experimental groups in randomized manner. Confounder effects were eliminated by ensuring that animals were placed in sterile cages in uniformly controlled environment along with supply of autoclaved food and water in institutional animal facility centre. The bedding material of all the cages were cleaned regularly and care was taken to remove the residual food pellets in timely manner.

### ‘The work has been reported in line with the ARRIVE guidelines 2.0’

***In-vivo* tumorigenicity assay:** Post co-culture, sorted CSCs (2x10^5^) of EC and 4T1 cells were inoculated into the mammary fat pads of female swiss albino and BALB/c mice respectively for development of solid tumors and to elucidate TEPs impact on the process.

### Experimental groups: CSC (control), CSC+RP, CSC+TEP for both EC and 4T1 models

Experimental unit: Cage of animals

Sample size: CSC (control) (n=6), CSC+RP (n=6), CSC+TEP (n=6); Total n=18, for both EC and 4T1 models. The sample size of n=6 for each group has been used as it provides unbiased 95% confidence limit.

Blinding: Animals were blinded by treating them identically.

Tumor growth was monitored twice a week and measured using Vernier callipers. Tumor area was calculated (length x width) and presented in mm^2^. Health of the animals was monitored daily. **Anaesthesia was not applied as an integral part of any invasive approach throughout the work on mice**. However, to dissect out the mature tumors (20mm size), the mice were euthanized through overdose of ketamine HCL (160 mg/kg) and xylazine (20mg/kg) according to the guidelines of Committee for the Purpose of Control and Supervision of Experiments on Animals. **Tumor burden was not exceeded beyond the size (1500-2000mm^3^) admissible by Institutional Animal Care and Ethics Committee.**

**Experimental metastasis model:** Post co-culture, sorted BCSCs (2x10^5^) of 4T1 and EC cells were injected through the tail veins of female BALB/c and swiss albino mice respectively for development of lung metastasis. The animals were monitored for 4 weeks after which they were euthanized by overdose of ketamine HCL (160 mg/kg) and xylazine (20mg/kg) according to the guidelines of Committee for the Purpose of Control and Supervision of Experiments on Animals to dissect out the organs. **Anaesthetics were not used for any invasive procedure throughout the animal experiments.** Lung metastasis was confirmed by macroscopic imaging of the nodules in murine lungs along with HE staining of the cryo-cut lungs sections.

### Experimental groups: CSC (control), CSC+RP, CSC+TEP for both EC and 4T1 models

Sample size: CSC (control) (n=6), CSC+RP (n=6), CSC+TEP (n=6); Total n=18, for both EC and 4T1 models. The sample size of n=6 for each group has been used as it provides 95% confidence limit.

### Experimental unit: Cage of animals

**Processing of tumors:** Procured tumors were digested with 1% collagenase for generating single cell suspensions. The cells were then washed with 1X PBS and were then used for various *in-vitro* assays.

**Cryo-sectioning:** Freshly procured tumor samples were fixed in 4% paraformaldehyde for 2 hours at room temperature followed by incubation at 4^°^C overnight in 30% sucrose. They were then snap chilled in liquid nitrogen and stored in -80 ^°^C for later use.

For cryo-sectioning, the frozen tissue samples were embedded in OCT (optimal cutting temperature compound, Leica biosystems, Wetzlar, Germany) and cut into 5µm sections using cryostat (Leica CM1950, Wetzlar, Germany). The sections were collected on poly-L-lysine coated slides and stored at -80^°^C for further use.

**Histology and HE staining:** The cryo cut tissue sections were stained with hematoxylin-eosin (HE) following standard staining protocol.

**Soft agar colony formation assay:** Briefly 5x10^3^ BCSCs, post co-culture (1:100; BCSC: platelet, 24 hours), were rested on upper 0.35% soft agar layer along with 1mL culture media. Untreated BCSCs were kept as control. This layer was placed upon botton agar bed of 0.7% agar and cell culture medium. The culture setup was maintained for 21 days with fresh media supplementation on every third day. The colonies were finally micrographed and the number of colonies was quantified.

**Contact independent transwell assay:** For transwell assay, magnetically sorted BCSCs were co-cultured in presence of 0.4µm transwell membrane (Hi-Media, Mumbai, India) in 1:100 (BCSC: platelet) for 24 hours. Following the incubation period, the transwell was removed and the cells were undertaken for further analysis.

**Wound healing assay:** Post co-culture, sorted BCSCs of MCF-7 and MDA-MB-23: were grown in 6 well plates until completely confluent. A scratch or wound was drawn using a cell scratcher and wound healing was observed by taking micrographs at different time points for 24 hours. Percentage wound closure was calculated as final area/initial area X 100%

**Matrigel invasion assay:** Post co-culture, sorted BCSCs (1x10^4^ cells) were serum starved for 3 hours after which they were layered on matrigel coated transwell inserts (8µm) (Corning-354480, New York, USA). These inserts with the cell suspension were placed in 24 well plate containing FBS as chemoattractant. This entire setup was maintained for 24 hours following which the invaded cells were fixed with paraformaldehyde and permeabilized by absolute methanol followed by staining with 0.2% crystal violet. Migrated cells from five random fields were photographed and quantified.

**Vascular mimicry (VM) assay:** For VM assay, 96 well flat bottom plates were coated with 60µL of growth factor reduced basement membrane (Matrigel; R&D System, MN, USA) and incubated at 37^°^C for solidifying. Roughly 5x10^3^ BCSCs in 150 µL of complete media were rested on the plates and incubated at 37^°^C for 24 hours. Following the incubation, the plates were micrographed and number of interconnected tubes per field was counted. Length and width of the tubes were quantified in ImageJ software.

**RT PCR:** Total RNA content of single cell suspensions was extracted by Trizol (Ambion, Thermo Fisher Scientific, MA, USA). cDNA was synthesized according to manufacturer’s protocol from it using Revert Aid First Strand cDNA Synthesis Kit (Thermo Fisher Scientific, MA, USA). Reverse transcriptase PCR was performed using 2X Go Taq Green Mix (Promega, WI, USA). Electrophoresis was done using 1.5% agarose gels and stained with ethidium bromide. PCR bands were visualized on ChemiDoc XRS+ (BioRad Laboratories, CA, USA), identified by image Lab softwareV5.1 and quantified using ImageJ software. List of gene specific primers used in PCR are given in Supplementary table T-2.

**Flow cytometry:** Briefly cells were stained with fluorescently tagged antigen specific antibodies and incubated in dark at 4^°^C for 30 minutes. For intracellular molecules, cells were treated simultaneously with 0.2% saponin (permeabilization buffer). Cells were finally washed, fixed with 1% paraformaldehyde and data was acquired using BD LSR Fortessa X-20 Cell Analyzer (Becton Dickinson, New Jersey, USA). For data collection Cell Quest Pro 5.1 and for analysis FlowJo (Becton Dickinson, New Jersey, USA) were used. Cell morphology was determined by FSC-A and SSC-A.

**Extraction of cytosolic and nuclear proteins:** Ice cold nuclear extraction buffer was added to the cell pellets and incubated at 4^°^C for 1 hour. The cells were centrifuged at 6000 RPM for 5 minutes and the supernatant was collected as cytosolic fraction. The pellet was dissolved in nuclear extraction buffer and vortexed for 30 minutes at 4^°^C. The samples were centrifuged at 12,000 RPM for 10 minutes. The supernatant thus obtained was the nuclear fraction.

**Extraction of total proteins and Western Blot:** Cells were lysed by incubating in RIPA buffer for 30 minutes at 4^°^C. This was followed by centrifugation for 30 minutes at 12,000 RPM at 4^°^C. Protein concentration of the lysates so obtained was determined by Bradford assay. 30-50µg of the protein lysates was separated on 12% SDS-PAGE and transferred onto nitrocellulose membrane using BioRad Gel Transfer system and bands were developed using ECL Kit (Advansta, CA, USA). Band intensity was quantified using Image Lab 6.2 software (Bio-Rad, California, USA).

**Immunofluorescence:** Targeted samples were harvested on poly-L-lysine coated glass slides and initially blocked with 5% BSA at RT. Cells were perforated with 0.15% Triton X-100 prior to blocking for staining intracellular molecules. Antigen specific primary antibodies were added to the section and incubated overnight at 4^°^C. Fluorescently tagged secondary antibodies were added next and incubated for 3 hours at RT. Finally, the slides were thoroughly washed and mounted with Fluoroshield DAPI 9Abcam, Cambridge, UK). Images were acquired using Olympus-BX53 microscope (Olympus Life sciences, Tokyo, Japan). Fluorescence intensity was evaluated using Image-J software and corrected total cell fluorescence (CTCF) was calculated using the formula CTCF= Integrated density- (Area of selected cell x Mean fluorescence intensity of background readings).

**Confocal Microscopy:** Platelet-BCSC interaction was visualized by confocal microscopy. Briefly, mammospheres were initially cultured in chamber slide. Binding of TEPs and RPs to mammospheres was allowed for another 24 hours. The cells were fixed in 2% PFA for 30 minutes, followed by perforation with 0.5% Triton X for 10 minutes. Nonspecific interaction was blocked by 5% BSA treatment. BCSCs were stained with fluorescently tagged PSGL1 and platelets with P-selectin and incubated in dark at 4^°^C overnight. Cells were washed with 1X PBS atleast thrice, mounted with DAPI and visualized with Olympus Fluoview FV3000. Images were analysed and quantified using ImageJ software.

**Immunohistochemistry:** Tissue sections were initially harvested on poly-L-lysine coated glass slides and kept in 1X PBS until the OCT was removed. Sections were treated with 3% H_2_O_2_ (Merck 17544) in methanol for 30 minutes, to block endogenous peroxidase. Non-specific sites were blocked by incubating the sections with 5% BSA at RT for 30 minutes followed by incubation with primary antibody overnight. After thorough washing with PBS-Tween 20, HRP tagged secondary antibody was added to the samples. AEC Substrate (Vector Laboratory SK4200) was used to develop chromogenic colour according to manufacturer’s protocol. Counterstaining was performed using Hematoxylin (Merck-HX68597049) for 40 seconds and then mounted with Vectamount (Vector Laboratories H5501). Image was acquired using Carl Zeiss Plan Achromat bright field microscope along with Axiocam 1058 color camera.

**siRNA mediated silencing *in-vitro*:** siRNA for human-WNT and human-VEGFR2 were constructed according to the manufacturer’s protocol *in vitro* using the Ambion Silencer^R^ siRNA construction kit (Life Technologies, USA). Gene specific siRNA and scramble control siRNA (Sigma-Aldrich) were added to *in vitro* setup to a final concentration of 50nM to the 2 hour-serum starved cells in presence of lipofectamine-2000 reagent (Invitrogen, USA). Sequence of primers utilized are mentioned below

***WNTsi Sense*:** 5’-AAGCAGGCTCTGGGCAGCTACCCTGTCTC-3’ ***WNTsi Antisense*:** 5’-AAGTACTGCCCAGAGCCTGCCCTGTCTC -3’ ***VEGFR2si Sense*:** 5’-AAGGTGCTGCTGGCCGTCGCCCCTGTCTC-3’

***VEGFR2si Antisense*:** 5’-AAGGCGACGGCCAGCAGCACCCCTGTCTC-3’

**Protein-protein interaction visualization:** String (Search Tool for the Retrieval of Interacting Genes/Proteins) database version 8.0 RRID: SCR_005223 was utilized to decipher the interactions between various proteins of interest. Based upon former reports of direct and indirect interactions, an interactome map illustrating this interrelationship between the proteins was generated. Each protein is assigned a colour and their interaction with the other proteins is represented by multicolour lines. The network properties include; nodes which represents the number of proteins in the interactome, edges depicting the number of interactions, followed by node degree which refers to the average number of interactions and finally clustering coefficient indicating the tendency of the network to form clusters.

**Statistical Significance:** Statistical significance was drawn from either Student *t*-test (for 2 groups) or one-way/two-way analysis of variance. Mean±SD of the results obtained from either three for *in-vivo* (n=6) or three-six for *in-vitro* (n=3-6) independent experiments were performed. To ensure normal distribution pattern, normality and log normality tests were performed. All data passed the Shapiro-Wilk test of normal distribution. Entire statistical analysis was performed using Graphpad Prism 8.4.2 software (Graphpad Software, San Diego, USA). Experimental results with p ≤ 0.05 have been considered as significant.

**Data availability:** Data supporting the present study will be available from the corresponding author upon reasonable request.

## Results

### Platelets of breast cancer patients differ morphologically and functionally from normal healthy platelets

Besides their longstanding role in wound healing, platelets crucial role in cancer is only modestly studied. To deeply elucidate their role in cancer progression and metastasis, we primarily investigated the differences between platelets of healthy donors and BC patients. For this, thrombocytes were isolated by double centrifugation method and characterized for their peculiarities in breast cancer (Fig 1a).

**Figure 1:**
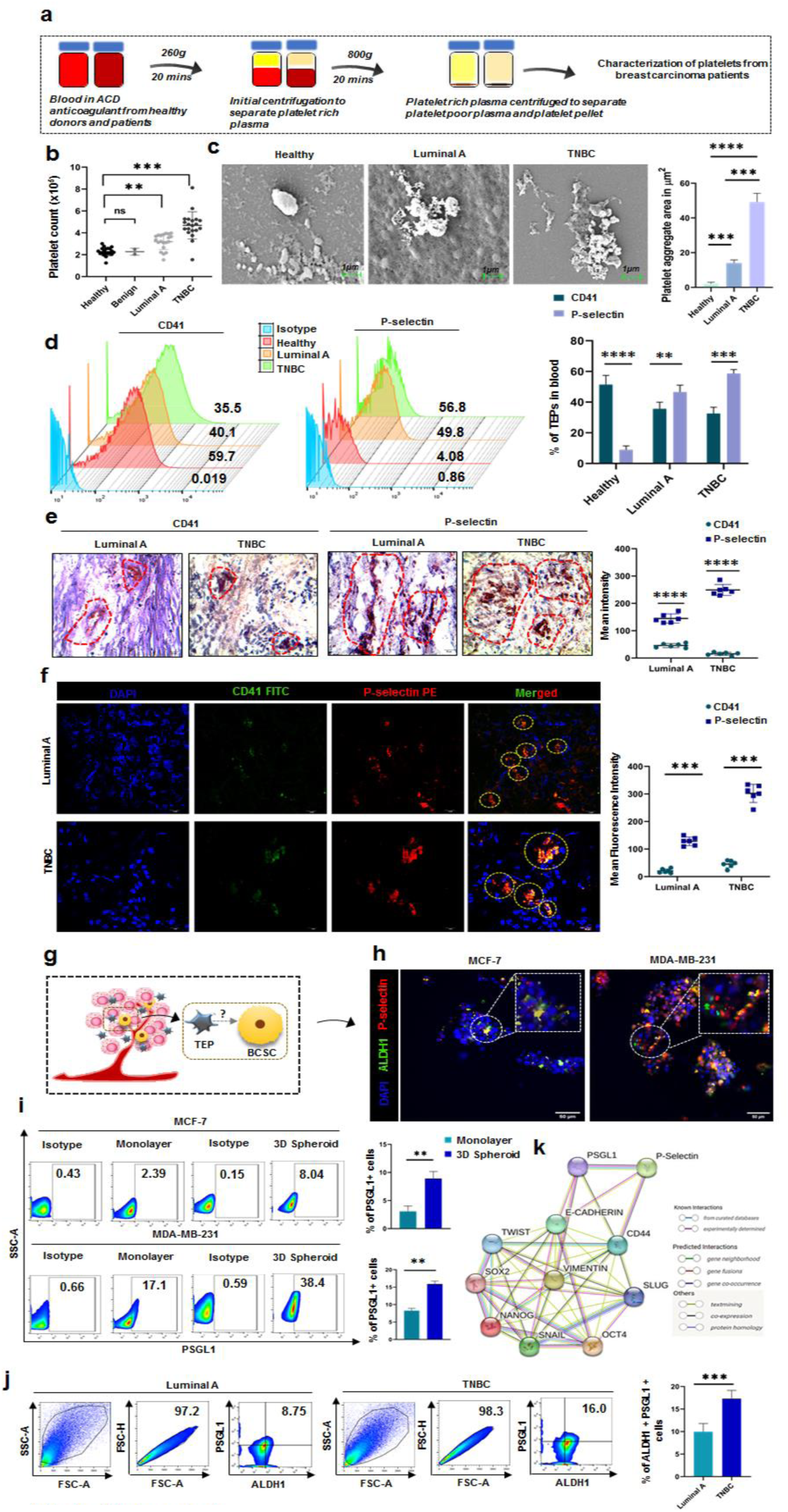
Platelets of breast cancer patients differ morphologically and functionally from normal healthy platelets: **a.** Schematic representation of the study. **b.** Representative scatter plot of platelet count in healthy (n=20), benign (n=2), luminal A(n=20), and TNBC (n=20) patients. **c.** SEM microscopy images of healthy, luminal A and TNBC patient’s platelets (scale 1µm). In bar-graph, (mean±SD) of platelet area in µm^2^ is provided. Statistical significance drawn from one-way ANOVA followed by Tukey’s multiple comparison test (n=6). **d.** Flow cytometric histogram plots showcasing frequencies of CD41, P-selectin, in healthy donors (n=11), luminal A (n=11), TNBC patients (n=11). Bar diagrams representing % of TEPs keeping (mean±SD) and statistical significance established from two-way ANOVA followed by Tukey’s multiple comparison test. **e**, **f.** Immunohistochemical micrographs at 40X magnification and confocal microscopy images at 100X magnification respectively of tumor sections from luminal A, TNBC stained with CD41-FITC and P-selectin-PE. Dotted lines representing positive zones are provided. Scatter plot (mean±SD) displays mean intensity with statistical significance inferred from two-way ANOVA followed by Tukey’s multiple comparison test. **g.** Schematic representation of probable interaction between TEP and CSC within the TME. **h.** Immunofluorescence micrographs at 40X magnification of mammospheres of MCF-7 and MDA-MB-231 stained with ALDH1-FITC and P-selectin-PE. Enlarged image of their interaction is provided in inset. **i.** Flow cytometric plots showcasing frequency of PSGL1 in monoloayer and 3D spheroids of MCF-7 and MDA-MB-231. In bar-graph, (mean±SD) of PSGL1% is provided and statistical significance is inferred from unpaired non-parametric t-test, followed by two-tailed p value. **j.** Flow cytometric plots showcasing frequency of ALDH1^+^PSGL1^+^ BCSCs in luminal A and TNBC tissue samples. Bar-graph represents (mean±SD) of PSGL1% is and statistical significance is inferred from unpaired non-parametric t-test, followed by two-tailed p value. **k.** Identification of the genes responsible for linking PSGL1 with stemness-metastasis using STRING database is given **b, c, d, e, f, i, j** *p<0.05, **p<0.01, ***p<0.001, ****p<0.0001, ns: not significant are indicated.

Retrospective study of patient records admitted at Chittaranjan National Cancer Institute, Kolkata, India, revealed that BC patients had platelet count >3 x10^5^/microL of blood (n=20 for healthy control, luminal A and TNBC patients).Further, upon comparison between luminal A and TNBC subtypes, it was observed that this augmentation in platelet count was more in TNBC (average platelet count ∼ 4.7x10^5^/ microL) than luminal A (average platelet count ∼ 3.1 x10^5^/ microL). Interestingly, in benign breast tumor patients (n=2), the mean platelet count was in the normal range (average platelet count of benign tumor patients ∼ 2.2 x 10^5^/ microL, average platelet count of healthy females ∼ 2.17 x10^5^/ microL). This further strengthened our observation of thrombocytosis due to underlying cancer condition (Fig 1b).

Next, morphological analysis revealed that, in comparison to healthy resting platelets which have a discoid appearance with approximately 1.5μm^2^ area, platelets of cancer patients were of altered morphology and formed huge aggregates. The average area of the aggregates in luminal A patients was ∼15 μm^2^, whereas in TNBC it was ∼45 μm^2^ (n=6). Additionally, they underwent cytoskeletal rearrangement producing numerous filopodia and lamellipodia (Fig 1c). The role of tumor cells in inducing platelet aggregation and activation is well established. Presence of such aggregates in the peripheral blood of BC patients confirmed the potential impact of tumor cells in this process. These platelets hence are termed as tumor educated platelets or TEPs

With the prevalence of thrombocytosis and cytoskeletal rearrangement in cancer, we next sought to investigate the functional alteration of these platelets. Following their isolation from peripheral blood by double centrifugation method from healthy donors and BC patients, flow cytometric analysis of P-selectin (activated platelet marker) and CD41(healthy platelet marker) was performed. In comparison to healthy normal (n=11), expression of P-selectin was elevated in both luminal A (n=11) and TNBC patients (n=11) than CD41. Further, amongst these subtypes, this upsurge was more prominent in TNBC than luminal A. On the other hand, normal healthy platelets were predominantly CD41^high^ (Fig 1d).

In addition to peripheral blood, status of platelets within the tumor microenvironment (TME), was also investigated by screening of breast tumor sections. Immunohistochemical analysis of CD41 and P-selectin stained micrographs showcased infiltration of CD41^low^P-selectin^high^ platelets within TME of both luminal A and TNBC patients, with higher infiltration in TNBC than luminal A (Fig 1e). This was further confirmed by confocal microscopy imaging that also demonstrated a similar trend of infiltration of CD41^low^P-selectin ^high^ platelets into the TME of both luminal A and TNBC subtypes (Fig 1f)

In an attempt to revert this activation of platelets, we treated TEPs with acetylsalicylic acid or aspirin as described earlier (Supplementary Figure S1a). Post blocking the platelet pellet was analysed for P-selectin expression by flow cytometry. Histogram analysis revealed that aspirin treatment could reduce P-selectin expression significantly in both the subtypes (Supplementary Figure S1b).

Given that TEPs are an integral component of the breast TME, we wanted to elaborate, if CSCs are a potential partner of TEPs, and if this alliance have any role in augmenting aggressiveness of the disease (Fig 1g). For this, lin^-^CD44^+^CD24^-^ BCSCs of MCF-7 and MDA-MB-231 where grown into mammospheres in 3D stem cell enrichment setup and co-cultured with TEPs. Following this, BCSCs were stained with ALDH1, which is a prominent CSC marker and TEPs with P-selectin. Immunofluorescence microscopy images confirmed the interaction between TEPs and BCSCs *in-vitro* (Fig 1h).

Further, we elucidated if BCSCs express PSGL1, the ligand of P-selectin. Pseudo-colour flow cytometric plots depicted that both MCF-7 and MDA-MB-231 tumor cells and their corresponding BCSCs express PSGL1. However, this expression was more pronounced in BCSCs than the tumor cells (Fig 1i). To prove this *in-vitro* observation *in-vivo*, single cell suspension of breast tumor tissue samples of luminal A and TNBC subtypes were analysed for the expression of PSGL1. In line with our *in-vitro* results, in *in-vivo* too, TNBC patients exhibited greater population of ALDH1^+^ PSGL1^+^ BCSCs than luminal A (Fig 1j). As PSGL1 was more prominent on BCSCs, we investigated if it has any role to play in metastasis and stemness. String analysis disclosed direct interaction of PSGL1 with metastasis marker VIMENTIN and breast CSC marker CD44 which further interacts with the other genes and transcription factors that are pivotal for the process (Fig 1k).

### TEP induces BCSC upregulation, governed by NANOG-OCT4-SOX2

Given the possible synergism of TEPs and BCSCs in disease advancement, their coaction was further analysed. We investigated whether this mutual association between TEPs and BCSCs have any role in augmenting the hallmarks of CSCs specifically for tumorigenicity, clonogenicity and MDR phenotype. For this, post co-culture, lin^-^CD44^+^CD24^-^ BCSCs were magnetically sorted from single cell suspensions of solid breast tumor samples of both luminal A and TNBC subtypes (Fig 2a). These cells were propagated in 3D-CSC-enrichment setup for 7 days for primary tumorsphere formation. A significant upsurge of tumorspheres count along with an increment in surface area was noted in presence of TEPs with respect to control CSCs in both luminal A and TNBC. Further, administration of RPs significantly lowered this surge, exhibiting lesser tumorsphere formation with reduced surface area (Fig 2b).

**Figure 2:**
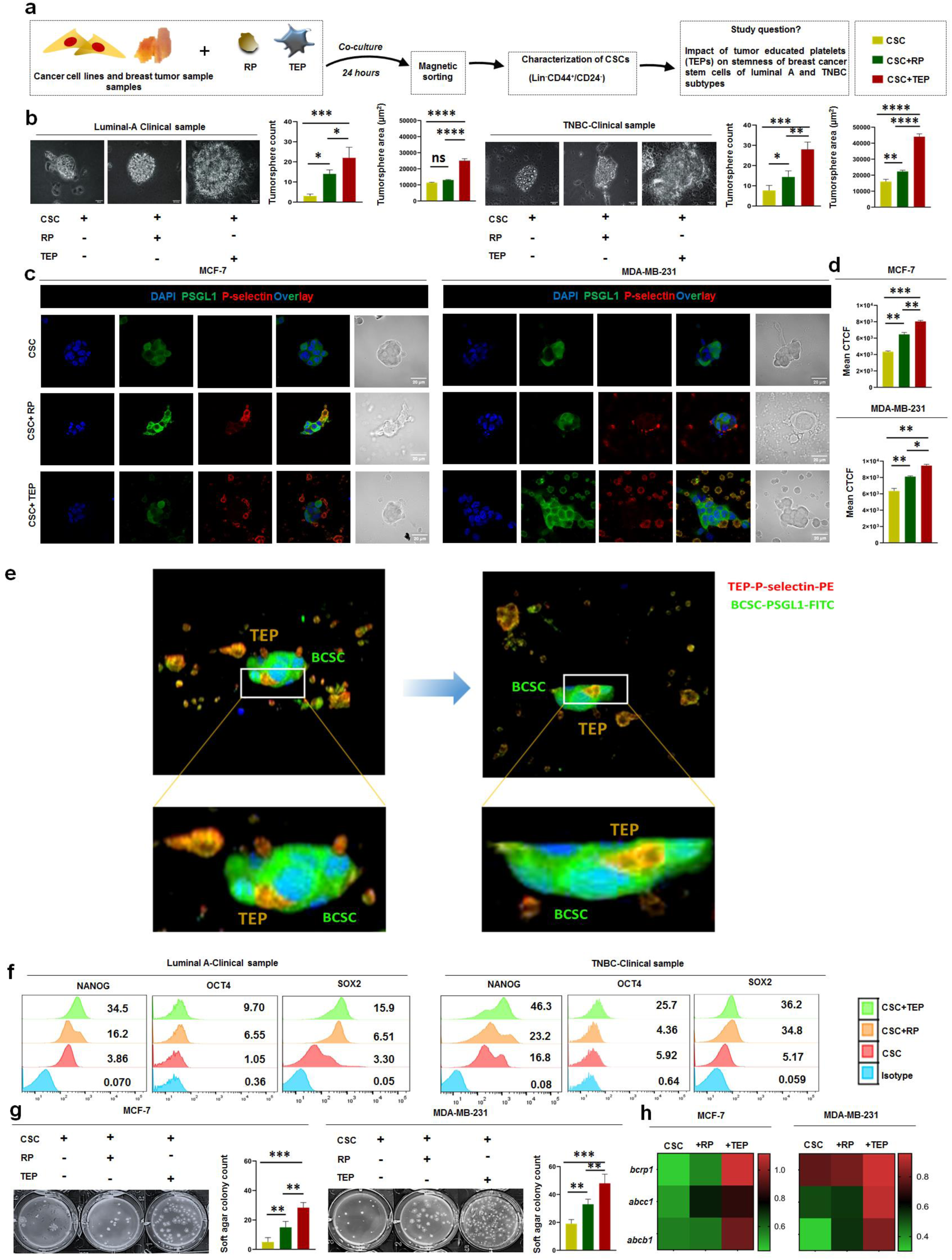
TEP induces BCSC upregulation, governed by NANOG-OCT4-SOX2: **a.** Schematic representation of experimental procedures and study question. **b.** Primary tumorspheres at 40X magnification from CSC, CSC+RP, CSC+TEP cohorts in luminal A and TNBC. In bar-graph, (mean±SD) for tumorsphere count and area is depicted (n=6). **c.** Confocal microscopy and bright field images at 100X magnification of mammospheres stained with PSGL1 (FITC) and TEP and RP stained with P-selectin (PE) of MCF-7 and MDA-MB-231 is presented. **d.** In bar-graph, (mean±SD) for corrected total cell fluorescence (CTCF) of the confocal images in MCF-7 and MDA-MB-231cohorts is depicted (n=3). **e.** Rotated composite view of the sphere positioned horizontally (left panel) displaying surface interaction between TEPs and BCSCs. Magnified view of the marked zone is displayed in the lower panel. A longitudinal section of the volume view of the field was cut to display the cell’s interior and rotated horizontally, illustrating infiltration of TEPs (right panel). Magnified view of the marked zone is displayed in the lower panel. **f.** Representative flow-cytometric histogram plots showcasing changes in the frequencies of NANOG, OCT4, SOX2 upon treatment with TEP and RP keeping non-treated CSC as control (n=6) in luminal A and TNBC clinical samples. *Inset*: blue, red, orange and green column denoting Isotype, CSC, CSC+RP, CSC+TEP respectively **g.** Soft agar colony images of CSC, CSC+RP, CSC+TEP of MCF-7 and MDA-MB-231 is represented. Number of colonies per field in each cohort is depicted in bar-graph (mean±SD) (n=3) **h.** Summary bar-diagram depicting mRNA expression of *bcrp1, abcc1, abcb1* keeping βactin as loading control in MCF-7 and MDA-MB-231 cohorts (n=3). All statistical significance inferred from one-way ANOVA followed by Tukey’s multiple comparison test. **b, d, g** *p<0.05, **p<0.01, ***p<0.001, ****p<0.0001, ns: not significant are indicated.

To investigate the changes in BCSC percentage post co-culture in 3D setup, cells were analysed via flow cytometry. It was revealed that there was a prominent increment in CD44^+^CD24^-^ BCSC-percentage in presence of TEPs with respect to controls and this uptrend was down-regulated upon treatment with RP in both the BC subtypes (Supplementary Figure S2a).

With the established role of TEPs in promoting mammosphere formation and upregulating CD44^+^/24^-^ BCSC population, we next sought to verify if the physical interaction between TEPs and BCSCs was mediated by P-selectin-PSGL1 axis. For this, lin^-^CD44^+^CD24^-^ BCSCs of MCF-7 and MDA-MB-231 were enriched in stem cell enrichment media to form mammospheres and co-cultured with TEPs and RPs, keeping non-treated mammospheres as control. Confocal microscopy imaging revealed that TEPs physically interacted with BCSCs in both luminal A and TNBC, and this interaction was facilitated by P-selectin and PSGL1 (Fig 2c, d, Additional file 2). Furthermore, it was noted that in RPs, the capacity of binding to BCSCs was considerably hampered, which in turn decreased the mean fluorescence intensity (Fig 2c, d). Interestingly, it was also observed that this interaction between TEPs and BCSCs was not just restricted to the surface, but TEPs were capable of infiltrating within the mammospheres, thereby depicting intra-cellular interactions (Fig 2e, Additional file 3).

Next, the status of transcription factors like NANOG, OCT4, SOX2 which orchestrate the various attributes of BCSCs were evaluated. A significant escalation in the expression of these molecules in presence of TEPs was noted from the flow cytometry analysis of the co-cultured samples of both the sub-types (Fig 2f).

One of the key attributes of CSCs is tumorigenicity, which is their ability to produce colonies from single cell. To delineate the possible role of TEPs in this process, soft agar colony formation assay was performed *in-vitro*. Post co-culture, magnetically sorted, BCSCs were plated on double layer soft agar bed for 21 days for colony formation. TEP influenced BCSCs showed increased colony formation compared to non-treated controls or RP influenced BCSCs in both MCF-7 and MDA-MB-231 (Fig 2g).

Resistance towards therapeutic agents make cancer treatment very much challenging. CSCs are wholly responsible for this therapy resistance. Analysis of MDR phenotype regulating genes via RT-PCR revealed increment in the expression of *bcrp1, abcc1* and *abcb1* in TEP influenced BCSCs of luminal A and TNBC thus affirming the potential role of TEPs in therapy resistance of cancer (Fig 2h).

Notably, the influence of TEPs on BCSCs was more pronounced in TNBC than luminal A. Cumulatively, these data suggests that TEPs physically interacts with the highly malicious BCSCs and thereby generate extremely tumorigenic, clonogenic, and therapy resistant BCSC sub-variants.

### TEP influenced BCSCs are highly invasive and metastatic in nature

The idea that CSCs are the foundation of metastasis is one of the main tenets of CSC model. Considering the fact that TEPs are able to generate highly malicious BCSC sub-variants and that this alliance indicated towards disease progression and additionally, CSCs being the basis of tumor invasion and metastasis, insights into the influence of TEPs on BCSC metastasis was elucidated. For this, following co-culture, magnetically sorted BCSCs were collected and evaluated for many key characteristics associated with the process (Fig 3a).

**Figure 3:**
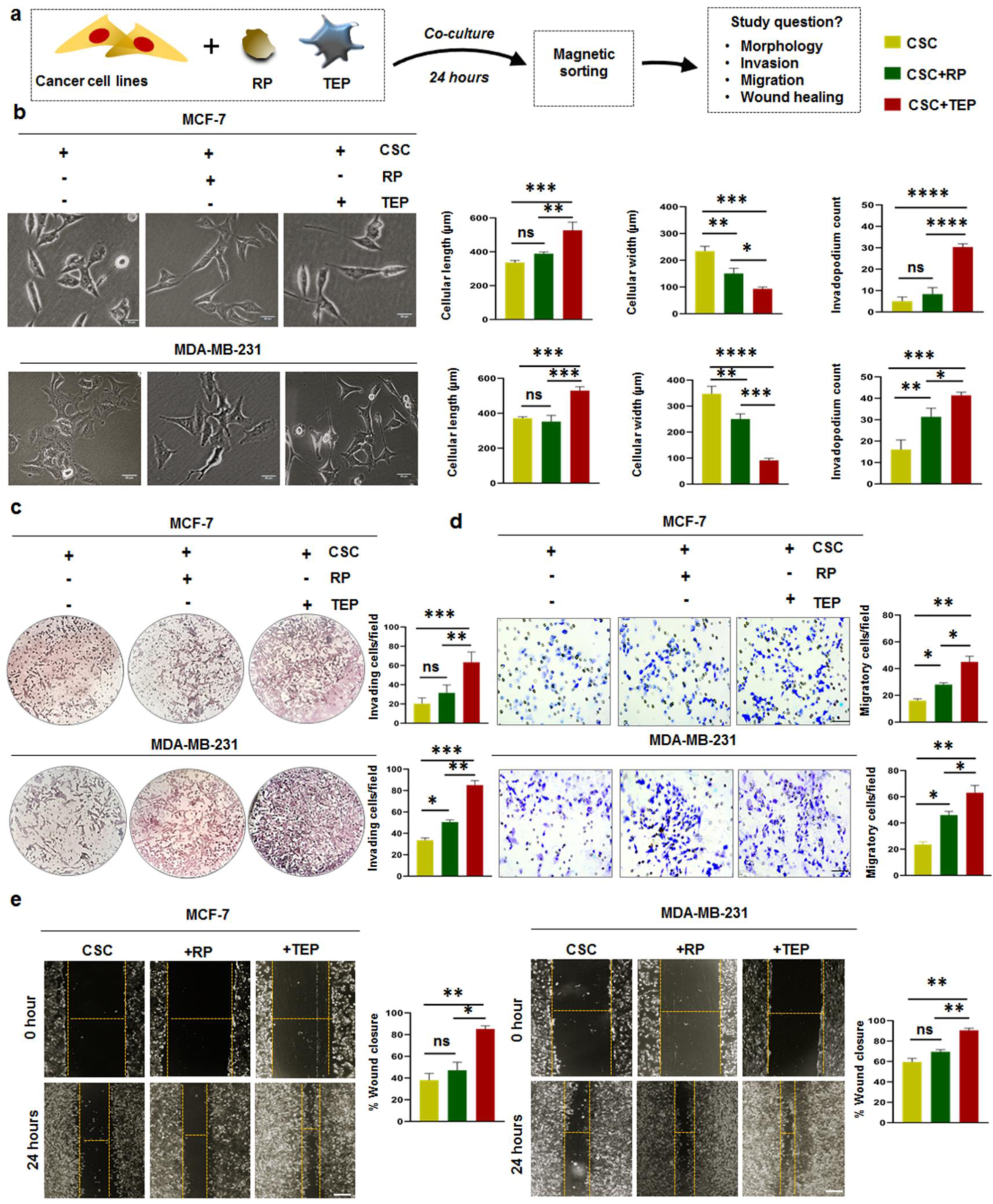
TEP influenced BCSCs are highly invasive and metastatic in nature: **a.** Schematic representation of isolating lin^-^CD44^+^CD24^-^ BCSCs by magnetic sorting from the co-culture setup of cancer cells with TEP and RP followed by illustration of the study questions. *Inset*: yellow, green and red column denoting CSC, CSC+RP, CSC+TEP. **b.** Representative micrographs at 40X magnification of post co-culture, sorted BCSCs of CSC, CSC+RP, CSC+TEP cohorts of MCF-7 and MDA-MB-231 showcasing distinct morphological features. Changes in cellular length and width and invadopodia count in CSC, CSC+RP, CSC+TEP of MCF-7 and MDA-MB-231 is presented in bar graph (mean±SD) with statistical significance drawn from one-way ANOVA followed by Tukey’s multiple comparison test (n=6). **c.** Micrographs of invading cells from matrigel invasion assay at 10X magnification of BCSCs of CSC, CSC+RP, CSC+TEP cohorts of MCF-7 and MDA-MB-231. In bar graph, invading cell count per field (mean±SD) is displayed and statistical significance inferred from one-way ANOVA followed by Tukey’s multiple comparison test (n=3). **d.** Representative images depicting migratory cells of transwell migration assay of CSC, CSC+RP, CSC+TEP of MCF-7 and MDA-MB-231. In bar graph, count of migratory cells per field (mean±SD) is given with statistical significance performed using one-way ANOVA followed by Tukey’s multiple comparison test (n=3). **e.** Representative micrographs at 10X magnification of wound healing assay at 0 hour and 24 hours across all three groups of MCF-7 and MDA-MB-231 is provided. Bar diagram (mean±SD) representing percentage wound closure in all the study groups of both the cohorts. One-way ANOVA followed by Tukey’s multiple comparison test was performed to draw statistical significance (n=3). **b, c, d, e,** *p<0.05, **p<0.01, ***p<0.001, ****p<0.0001, ns: not significant are indicated.

Morphological analysis through phase-contrast microscopy revealed that these sorted BCSCs (from both MCF-7 and MDA-MB-231) in the monolayer adopted a very linear architecture with increased cellular length and reduced cellular width under influence of TEPs. Conversely, in the presence of RP, these cells had almost similar cellular length like non-treated controls, but their width was reduced significantly in their comparison. Furthermore, TEP influenced BCSCs underwent cytoskeletal rearrangement producing numerous invadopodia on their surface. These structures facilitate focal degradation for invasion and metastasis (Fig 3b). Immunofluorescence staining of cofilin, an actin binding cytoskeletal protein revealed increased nuclear expression in TEP∼BCSCs than the control groups of MCF-7, suggesting its role in supporting the development of invadopodium (Supplementary Figure 3a).

This was further corroborated with the results of matrigel invasion assay, which showed that TEP-impacted BCSCs had an enhanced invasive potency than RP-influenced or control BCSCs, with higher number of invading cells per field in both MCF-7 and MDA-MB-231 (Fig 3c).

Additionally, it was discovered by transwell migration experiment that TEP-influenced BCSCs were also more mobile than controls, with a greater number of migratory cells per field in both the subtypes. Furthermore, administration of RPs reduced the migratory capacity of these invasive BCSCs (Fig 3d). The observations of wound healing assay provided more evidence for this. It was uncovered that TEPs elevated the extent of wound closure in these cells than controls (Fig 3e).

### TEP influenced BCSCs are skewed towards mesenchymal lineage

Now that the critical role of TEPs in facilitating invasion and metastasis of BCSCs has been demonstrated, we likewise attempted to clarify the roles played by transcription factors, proteins, chemokines and *mmps* in the process. In light of this, post co-culture magnetically sorted BCSCs were examined in order to decipher modifications in the expression of relevant molecules (Fig 4a).

**Figure 4:**
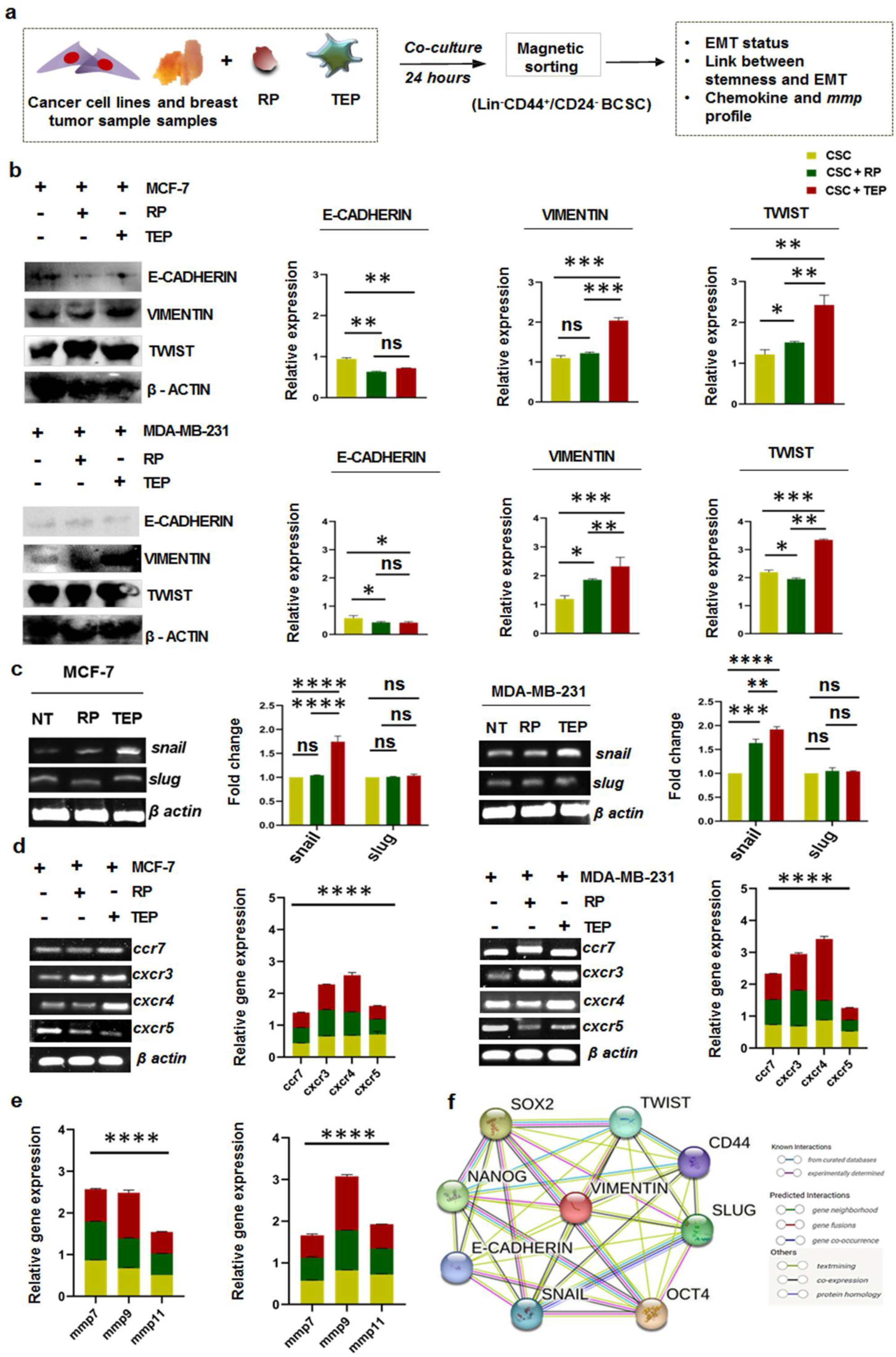
TEP influenced BCSCs are skewed towards mesenchymal lineage: **a.** Illustration of isolating lin^-^CD44^+^CD24^-^ BCSCs by magnetic sorting from the co-culture setup of cancer cells with TEP and RP followed by depiction of the study questions. **b.** Representative cropped western blot images of EMT and metastasis related proteins E-CADHERIN, VIMENTIN, TWIST keeping β-ACTIN as control in CSC, CSC+RP, CSC+TEP of MCF-7 and MDA-MB-231 is provided. Bar diagram (mean±SD) depicting relative protein expression is given. Two-way ANOVA followed by Tukey’s multiple comparison test was the source of statistical significance (n=3). *Inset*: yellow, green and red column denoting CSC, CSC+RP, CSC+TEP. Uncropped full-length blots with protein ladder is provided in Additional file 5. **c.** mRNA expression of genes *snail* and *slug* by RT-PCR, with β-ACTIN as control is presented in CSC, CSC+RP, CSC+TEP cohorts of MCF-7 and MDA-MB-231. In bar graph (mean±SD) quantified values of relative fold change is presented with statistical significance inferred from two-way ANOVA followed by Tukey’s multiple comparison test (n=3). Uncropped full-length gel is provided in Additional file 5. **d.** mRNA expression of chemokine related genes in CSC, CSC+RP, CSC+TEP of both the cell lines by RT-PCR is represented. Summary bar graphs of relative gene expression is given. Uncropped full-length gel is provided in Additional file 5. Statistical significance inferred from two-way ANOVA followed by Tukey’s multiple comparison (n=3). **e.** Representative summary bar-graphs of mRNA expression of matrix metalloproteinases (*mmp*7, 9, 11) CSC, CSC+RP, CSC+TEP cohorts of MCF-7 and MDA-MB-231. Two-way ANOVA followed by Tukey’s multiple comparison was performed to draw statistical significance (n=3). **f.** Identification of the genes responsible for linking stemness and metastasis using STRING database is provided. **b, c, d, e,** *p<0.05, **p<0.01, ***p<0.001, ****p<0.0001, ns: not significant are indicated.

Western blot analysis of EMT-related transcription factors revealed that extent of upsurge in the expression of VIMENTIN, TWIST and reduction in expression of E-CADHERIN was more in TEP treated BCSCs than controls in both MCF-7 and MDA-MB-231, confirming their mesenchymal lineage (Fig 4b). This was further corroborated by the results of RT-PCR analysis of *snail* and *slug* which demonstrated elevation in the expression of *snail* in BCSCs upon TEP treatment, validating their mesenchymal predisposition. Nevertheless, *slug* remained at a steady level throughout the groups (Fig 4c).

Next, a panel of chemotaxis-associated genes was evaluated for expression status using RT-PCR in order to determine the likely migration sites of these invasive, metastatic TEP-influenced BCSCs. The TEP-treated group exhibited increased expression of *cxcr4* and *mmp9* amongst these molecules (Fig 4d, e). Both *cxcr4* and *mmp9* have been reported to play pivotal role in regulating cancer metastasis (24–27).

Finally, to ascertain the link between stemness and metastasis, string data base was utilized which revealed the interactions between the various proteins that have been reported to be involved in interconnecting stemness with metastasis (Fig 4f).

Considering TEPs significant contribution in BCSC aided disease progression, we next investigated whether such influence was mediated by soluble factors (contact independent) or receptor-ligand interaction dependent (cell-cell contact dependent) mechanisms. For this, lin^-^ CD44^+^CD24^-^ BCSCs were magnetically sorted from single cell suspensions of solid breast tumor samples of both luminal A and TNBC subtypes post co-culture with TEP and RP, keeping non-treated BCSCs as control. For contact dependent interaction, BCSCs and TEPs were cultured together in the same well, whereas in the other setup they were physically separated using 0.4µm transwell inserts. This ensured that the only means of communication between BCSCs and TEPs was through soluble cellular secretions (Supplementary Figure S4a). In both luminal A and TNBC, TEP-induced BCSC were skewed towards the mesenchymal state predominantly in the contact dependent setup as indicated by the strong expression of VIMENTIN and low expression of E-CADHERIN (Supplementary Figure S4b).

*In-vivo*, luminal A and TNBC breast carcinoma tissue sections were examined for any indication of physical contact-dependent interaction between TEPs and BCSCs in order to further corroborate the *in-vitro* results. For this, BCSCs were tagged with stem cell marker OCT4 and TEPs with P-selectin (Supplementary Figure S4c). Immunofluorescence microscopy images revealed their proximity, thereby, validating their contact-dependent interaction. Their co-localization was affirmed by Mander’s co-localization coefficient (luminal A; M1-0.907 and M2-0.580) (TNBC; M1-0.958 and M2-0.657)

### TEP influenced BCSCs are highly tumorigenic, promotes angiogenesis and induces lung metastasis *in-vivo*

In context with the fact that TEPs are playing crucial role in enhancing stemness, clonogenicity of CSCs and their alliance with these cells elevated the invasiveness, migration, and metastasis of CSCs of BC subtypes *in-vitro,* we further ought to corroborate these findings *in-vivo* (Fig 5a).

**Figure 5:**
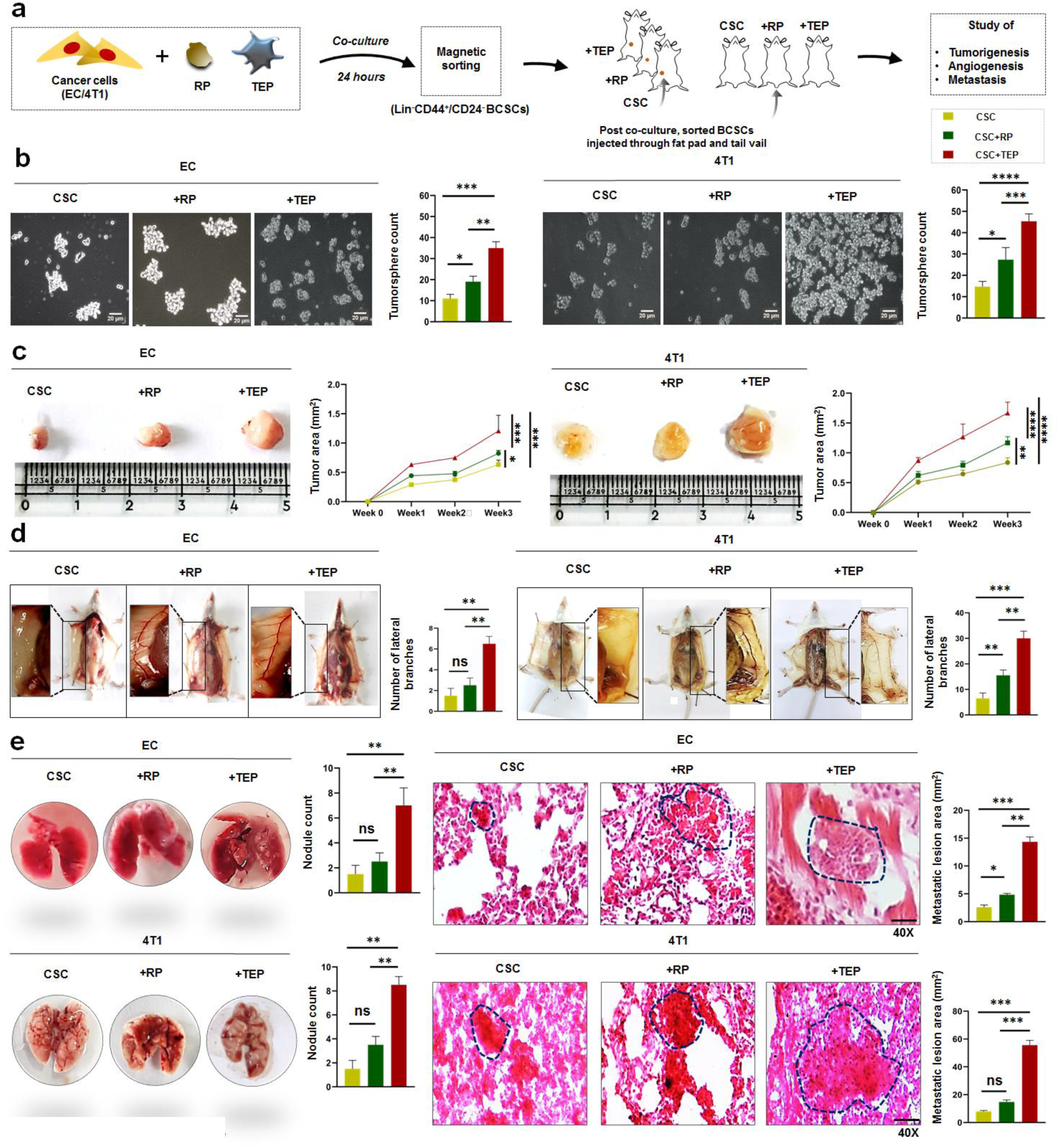
TEP influenced BCSCs are highly tumorigenic, promotes angiogenesis and induces lung metastasis *in-vivo*: **a.** Diagrammatic representation of magnetic sorting for isolation of lin^-^CD44^+^CD24^-^ BCSCs from co-culture setup of cancer cells (EC, 4T1) with TEP and RP and inoculation of CSC, CSC+RP, CSC+TEP into the mammary fat pads and tail vein of mice. *Inset*: yellow, green and red column denoting CSC, CSC+RP, CSC+TEP. **b.** Representative images at 10X magnification of primary tumorspheres in CSC, CSC+RP, CSC+TEP cohorts of EC and 4T1 cell lines. In bar-graph, mean±SD for tumorsphere count is displayed. Statistical significance is inferred from one-way ANOVA followed by Tukey’s multiple comparison test (n=6). **c.** *In-vivo,* BCSCs isolated from the co-culture of EC +/- TEP/RP and 4T1+/- TEP/RP was subcutaneously injected into the mammary fat pad of female swiss mice and BALB/c mice respectively for development of primary tumor. Representative photographs of harvested tumors with scale are presented (n=6). Tumor growth curve depicting tumor area in mm^2^ (mean±SD) at each time point is exhibited. **d.** Illustrative photographs of tumor draining blood vessel with lateral branches depicting angiogenesis across all groups of EC and 4T1 is presented (n=6). In bar-diagram, mean±SD of lateral branch count is given. Statistical significance drawn from one-way ANOVA followed by Tukey’s multiple comparison test (n=6). **e.** Photographs of lung nodules across the groups of EC and 4T1 is shown. Bar graphs (mean±SD) representing nodule count is given. Statistical significance (mean±SD) inferred from one-way ANOVA followed by Tukey’s multiple comparison test (n=6). Micrographs of Hematoxylin-Eosin stained lung tissue sections at 40X magnification depicting metastatic deposition in CSC, CSC+RP, CSC+TEP of EC and 4T1 is presented. Dotted lines depicting boundaries of deposition is given. In bar-graph, mean±SD for metastatic area in mm^2^ is presented and statistical significance drawn from one-way ANOVA followed by Tukey’s multiple comparison test (n=6). **b, c, d, e,** *p<0.05, **p<0.01, ***p<0.001, ****p<0.0001, ns: not significant are indicated.

BCSCs isolated from the co-culture setup and enriched in 3D stem cell enrichment media, elaborated increased tumorsphere formation in both the less aggressive EC cells as well as in more aggressive 4T1(Fig 5b).

In an effort to explore the influence of TEP-BCSCs on tumorigenesis *in-vivo*, BCSCs were isolated from the co-culture setup of EC and 4T1 with TEP and RP and 2x10^5^ cells were injected into the mammary fat pad of female swiss and BALB/c mice respectively to initiate primary tumor growth. Compared to BCSCs treated with RP or non-treated controls, tumor growth expanded dramatically in mice injected with TEP-BCSCs (Fig 5c).

The development and progression of tumor also requires uninterrupted supply of nutrients and oxygen to the rapidly proliferating cells. This is achieved either by neovascularization or by angiogenesis and also by vascular mimicry. From the primary tumor experiment, it was also observed that TEP-BCSC group had thicker vasculature with numerous lateral branching draining towards the tumor in comparison to RP-BCSC in both EC and 4T1 setup. Conversely, the thickness, and number of lateral branches from the primary blood vessel were all dramatically reduced in the non-treated control group (Fig 5d). Thus, TEPs ensured the steady supply of oxygen and nutrients to the tumor cells, which was significantly impeded in RP influenced cohort.

To further elaborate TEPs role in these events, vascular mimicry was performed *in-vitro*. For this, post co-culture sorted BCSCs were rested on matrigel and allowed to form tubes for 24 hours. TEP treated BCSCs in both MCF-7 and MDA-MB-231 showed increment in tube number with an upsurge in tube length than control CSCs or RP-CSCs which produced fewer tubes with reduced dimensions across the cell lines (Supplementary Figure S5a).

Moreover, we sought to determine if TEP-BCSCs may induce metastasis without regards to the primary tumor growth. In light of this, experimental metastasis was performed. To accomplish this, 2x10^5^ BCSCs were intravenously injected into the tail veins of female swiss and BALB/c mice, which were isolated from the co-culture setup of EC-TEP/RP and 4T1-TEP/RP respectively. A greater number of tumor nodules, with increased surface area of the metastatic foci was observed from macroscopic imaging as well as HE staining of the affected lungs, in both the murine models of TEP influenced BCSC cohort in comparison to the controls (Fig 5e). No significant changes were observed in the morphology and weight of liver, spleen and lymph nodes of these mice (Supplementary Figure 6a, b).

### TEP influenced stemness and metastasis of BCSCs is mediated in the downstream by WNT-β-Catenin-VEGF-VEGFR2 cascade

Considering the fact that physical contact dependency is involved in TEP-mediated stemness and BCSC metastasis, a number of probable receptor-ligand signaling pathways were investigated for potential relevance (Fig 6a).

**Figure 6:**
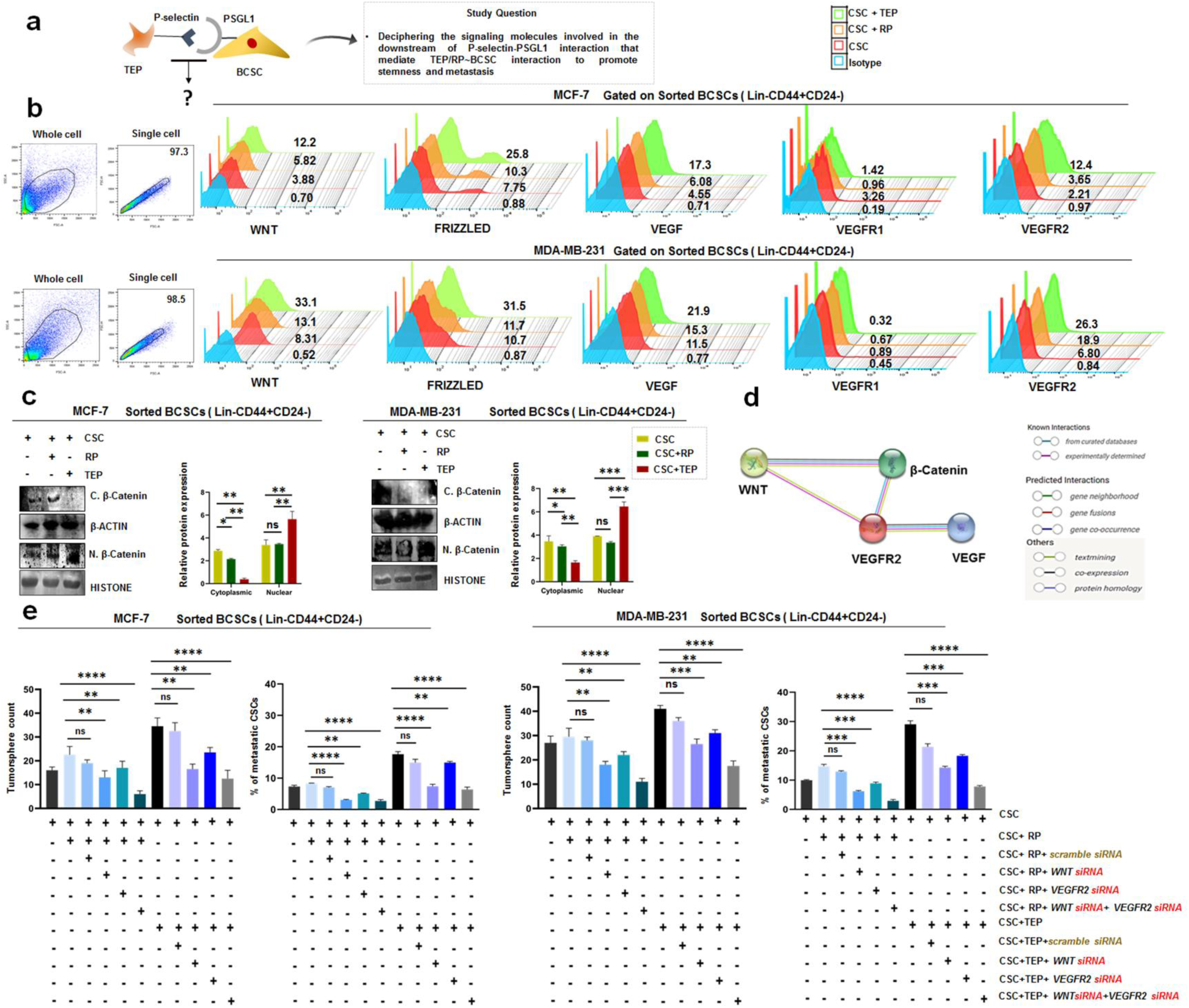
TEP influenced stemness and metastasis of BCSCs is mediated by WNT-β-Catenin-VEGF-VEGFR2 cascade: **a.** Schematic representation of receptor-ligand interaction between BCSCs and TEP/RP. **b.** Representative flow-cytometric histograms depicting expression of proteins WNT, FRIZZLED, VEGF, VEGFR1 and VEGFR2 on the surface of CSC, CSC+RP, CSC+TEP of MCF-7 and MDA-MB-231. *Inset*: blue, red, orange and green column denoting Isotype, CSC, CSC+RP, CSC+TEP respectively **c.** Illustrative western blots for cytoplasmic and nuclear fractionation of β-Catenin keeping β-ACTIN and HISTONE as respective controls across all the groups in both MCF-7 and MDA-MB-231. Uncropped full-length blots with protein ladder is provided in Additional file 5. Bar diagram (mean±SD) depicting relative protein expression is presented. Statistical significance inferred from one-way ANOVA followed by Tukey’s multiple comparison test (n=3). *Inset*: yellow, green and red column denoting CSC, CSC+RP, CSC+TEP. **d.** String analysis illustrating link between WNT-β-Catenin-VEGF-VEGFR2 is provided. **e.** Representative bar graph (mean±SD) depicting tumorsphere count and % of metastatic BCSC (CD44^+^24^-^VIMENTIN^+^E-CADHERIN^-^) across all the experimental groups of MCF-7 and MDA-MB-231 in presence and absence of WNT siRNA and VEGFR2 siRNA is provided. Statistical significance inferred from one-way ANOVA followed by Tukey’s multiple comparison test (n=4). **c, e** *p<0.05, **p<0.01, ***p<0.001, ****p<0.0001, ns: not significant are indicated.

CSCs were initially screened using RT-PCR for the conventional pathways governing stemness and metastasis. The results demonstrated a boost in *wnt* expression but no changes in *notch1* or *notch 4* in the TEP-BCSC cohorts of both MCF-7 and MDA-MB-231 (Supplementary Figure S7a). At the protein level too, this elevation in the expression of WNT and its receptor FRIZZLED was observed (Fig 6b). The cytoplasmic to nuclear translocation of β-Catenin further reinforced our findings (Fig 6c). Moreover, we discovered multiple studies elucidating the role of VEGF in controlling stemness and metastasis of cancer cells. In accordance with this, BCSC’s *vegf* and surface *vegf* receptor status was examined. Compared to the non-treated control CSCs, the TEP influenced cohort had higher mRNA expression of *vegf* and its corresponding receptor *vegfr2*. Nonetheless, *vegfr1* remained almost unchanged throughout the groups (Supplementary Figure S7a). Similar observation was also noted at the protein level (Fig 6b). Additionally, string analysis strengthened our observation by disclosing direct link between the targeted molecules (Fig 6d). Taken together, these findings confirmed that the influence of TEPs on BCSCs is mediated via WNT-β-Catenin-VEGF-VEGFR2 in the downstream to promote stemness and metastasis in BC subtypes.

To further verify the impact of this cascade, WNT and VEGFR2 were knocked out both individually as well as concomitantly *in-vitro* by siRNA in BCSCs of both MCF-7 and MDAMB-231. Following the incubation period, co-culturing was executed using TEP and RP and cells were propagated in 3D CSC enrichment media for formation of primary tumorspheres (Fig 6e, Supplementary Figure S8a). Inhibition of WNT and VEGFR2 affected tumorsphere count significantly, but the greatest reduction was observed when both WNT and VEGFR2 were targeted together. Surprisingly, in RP-BCSC cohort with dual knockdown of WNT and VEGFR2 exhibited the highest reduction in tumorsphere count amongst all the experimental groups in both MCF-7 and MDA-MB-231. A similar trend was also noted in the percentage of metastatic CSCs (CD44^+^24^-^VIMENTIN^+^E-CADHERIN^-^) of both the sub-types (Fig 6e, Supplementary Figure S8b). Cumulatively, our findings validate the involvement of WNT-β-Catenin-VEGF-VEGFR2 signaling pathway in the downstream of TEP-P-selectin-BCSC-PSGL1 interaction to mediate stemness, metastasis and overall aggressiveness of BC subtypes.

## Discussion

Pro-metastatic role of platelets has been reported in several malignancies including ovarian, myeloma, colon and Lewis lung carcinoma (28). In this current study, we describe how platelets contribute to the progression of BC in two subtypes: luminal A (the less aggressive type) and TNBC (the most aggressive type). Thrombocytosis was prevalent in both of these variants and we uncovered that a higher proportion of P-selectin^+^ platelets were present in the peripheral blood of BC hosts. Analysis of their medical records revealed that these patients did not suffer from any other health conditions at the time of diagnosis that could boost the production of platelets. This substantiated the existence of thrombocytosis and eliminated the probability of thrombocythemia.

Platelet’s α-granules are a repository of P-selectin. They degranulate after activation, revealing P-selectin on their surface. This suggested that platelets in these two BC subgroups were activated. Furthermore, TNBC had an elevated percentage of activated TEPs than luminal A, which indicated towards the potential involvement of TEPs in disease aggressiveness. Moreover, platelets of cancer patients formed huge aggregates and had an elaborate system of filopodia and lamellipodia, reinforcing their activated state. Our results are consistent with preliminary reports on murine cancer models, where it has been demonstrated that excessively metastatic tumors promote platelet activation (29). In line with previous reports on, the interaction of platelets with tumor cells to generate tumor educated platelets or TEPs (30–31), we demonstrate that TEPs can infiltrate into the tumor and are an indispensable component of the breast TME. Moreover, TNBC patients exhibited higher TEP infiltration than luminal A. This further strengthened the possible contribution of TEPs in promoting cancer aggression.

We next sought to ascertain TEPs functionality within TME. We discovered that TEPs and BCSCs remained in close proximity inside TME, suggesting the likelihood of their interaction. Additionally, TNBC∼TEPs demonstrated a stronger affinity towards BCSCs than luminal A. This propinquity of TEPs towards BCSCs, especially in TNBC, confirmed that TEPs tend to form an alliance with BCSCs, that dictates the overall disease outcome.

Further, analysis of the surface receptors revealed that BCSCs express PSGL1 which binds with TEPs P-selectin and this kinship facilitates tumor progression. PSGL1 is reported to have significant role in metastasis (32). In prostate and small cell lung cancer, malignant cells with high PSGL1 expression is reported to be capable of distant metastasis (32–34).

Upon in-depth investigation of this TEP and BCSC interplay *in-vitro*, it was found that TEPs elevated BCSC population, generating extremely tumorigenenic and clonogenic sub-variants. Analysis of transcription factor that regulate various attributes of CSCs disclosed that under the influence of TEPs there was significant upregulation of NANOG, OCT4 and SOX2. Also, these TEP-BCSCs overexpressed genes related to MDR phenotype such as *abcc1, abcb1 and bcrp1*. The role of TEPs in drug resistance has already been established in various tumors (35–36). Our investigation thus strengthened TEPs role in therapy resistance and disease relapse which are characteristic hallmarks of CSCs.

The precise role of TEPs on metastasis of BC was delineated next. It was found that TEP influenced BCSCs remained in a typical mesenchymal state. The link between EMT and stemness has been demonstrated in a number of earlier studies, wherein EMT has been connected to the development and maintenance of CSCs, conferring them with stemness and flexibility (37–38). TEP-BCSCs also attained an elongated architecture with numerous invadopodia on their surface. This invasive morphology strongly supported their mesenchymal nature. Matrigel invasion assay, migration assay and wound healing assay further stiffened the metastatic nature of TEP influenced BCSCs. Overexpression of VIMENTIN, TWIST, SNAIL and downregulation of E-CADHERIN complemented our observations.

Elevation of NANOG and OCT4 might play multi-faceted role by maintaining stemness and at the same time promote invasion by downregulating E-CADHERIN (39). Furthermore, by upregulating the expression of VIMENTIN and suppressing E-CADHERIN, increased TWIST expression may promote EMT (40). Additionally, through its interaction with NANOG, it may enhance stemness and resistance to drugs by augmenting the expression of *bcrp1*(40).

With the inception of TEPs function in facilitating metastasis, the secondary site of migratory BCSCs was delineated. Elevated *cxcr4* has been linked to BC metastasis to liver, brain, lungs, and lymph nodes according to literature reports (24–27). Furthermore, because of its great capacity for ECM breakdown, *mmp9* plays a critical role in metastasis and its high expression has been correlated with poor prognosis in BC, according to clinical studies (24–27).

Experimental metastasis in both female swiss and BALB/c murine models revealed that these BCSCs infiltrate through the blood vessels to initiate distant pulmonary metastasis. Our results also were supported by a similar observation where it was reported that in patients with osteosarcoma, high CXCR4 and MMP9 expression promoted lung metastasis (24).

With the established role of TEPs in promoting stemness and metastasis of BCSCs via physical contact, we next explored the downstream mediators of P-selectin-PSGL1 receptor ligand interactions. Interestingly, we observed that the WNT-β-Catenin and VEGF-VEGFR2 axis were elevated among the many signaling cascade components involved in controlling stem cell fate. Previous reports have mentioned about the collaborative role of WNT and NOTCH to control stemness, and that their patterns of expression are analogous (41). But we report a non-conventional signaling in which WNT-β-Catenin forms alliance with VEGF-VEGFR2 instead of NOTCH in TEP-BCSC interaction. The role of VEGF-VEGFR2 cascade in promoting stemness and metastasis of BCSCs has already been established (41–42). Our observation thus correlates with previous reports of crosstalk between WNT and VEGF in breast cancer (42). Also, the direct role of β-Catenin in inducing VEGF production is very well reported in malignancies like colorectal cancer (43–46).

Finally, the therapeutic role of aspirin was explored. Novel anti-platelet agent aspirin is well established to block platelet activity and impede its procarcinogenic effect (47–50). This is in line with our results, where aspirin reduced the activation of TEPs in BC subtypes. Aspirin treatment declined P-selectin expression on TEPs reverting them almost to RP state. This blocked its interaction with PSGL1, thereby hindering the complete downstream signaling cascade, ultimately reducing stemness, tumorigenicity, MDR phenotype and metastasis of BCSCs. The most interesting observation was noted in the RP-BCSC cohort, with the knockdown of both WNT and VEGFR2. The greatest reduction in tumorsphere count and metastasis was recorded in this group. A substantial explanation could be that blocking P-selectin hampered the receptor-ligand interaction, and further dual knockdown of WNT-VEGFR2 provided additional stress. This proves the direct link between P-selectin and WNT-VEGFR2 cascade

Thus, this study discloses the importance of tiny anucleated platelets in malignant disease progression and aggressiveness. Their relevance as therapeutic targets are supported by their capacity to interact with the vicious CSCs to produce extremely aggressive and metastatic sub-variants via EMT. Besides, their significance in fostering disease aggression is aided by the fact that they interact with TNBC to a larger extent than luminal A. Anti-platelet medications may thus offer an innovative approach in BC treatment. Neutralization of the WNT-VEGFR2 axis may potentially be investigated in conjunction with it. Further, impact of TEPs on the other subtypes of BC may also be elucidated.

## Conclusions

In summary, in search of other cellular partners that tumor educated platelets may interact with to promote metastasis in BC, we found cancer stem cells as the putative target of these platelets. We initially deciphered that, platelets in BC patients tend to form aggregates and undergo cytoskeletal rearrangement. Further, we came to know that, TEPs have high expression of P-selectin than healthy platelets, confirming the degranulation of α-granules, which is a repertoire of P-selectin. As platelets degranulate under activation stimuli, it suggested that platelets are in activated TEP state in BC patients. Not only did we observe the presence of such P-selectin ^high^ TEPs in the peripheral blood of luminal A and TNBC BC patients, but such TEPs were also an integral component of breast TME. Additionally, it was also noted that such population of TEPs were higher in TNBC than luminal A. This suggested that TEPs may have an important role in promoting aggressiveness of the disease.

We next screened BCSCs for the expression of PSGL1, which is the ligand of P-selectin. In line with previous reports that have mentioned that tumor cells express PSGL1 (51), we too observed that breast tumor cells are PSGL1^+^, but interestingly, in comparison to the tumor cells, PSGL1 remained elevated in BCSCs of both luminal A and TNBC subtypes. Substantially, we deciphered that, TEPs via P-selectin receptor binds to PSGL1 ligand on the surface of BCSCs. As a result, an intracellular signal is initiated within the BCSCs. This signal stabilizes β-Catenin, which translocate into BCSC nuclei, and enhances transcription of NANOG, OCT4, SOX2 along with TWIST, SNAIL and VIMENTIN. Simultaneously, it stimulates autocrine VEGF production which binds to VEGFR2 receptor on surface of BCSCs. Together, the signaling cascade intensifies stemness and directs BCSCs towards metastatic phenotype via EMT. Further, suppression of P-selectin expression by administration of aspirin, reduced the overall intensity of the scenario by regressing stemness and metastasis.

This study may pioneer in connecting two potent players of metastasis and impact of their alliance in promoting tumor progression, by linking stemness with EMT and metastasis. This would help to explain the observed physiological potential of TEPs in cancer in the upcoming future and provide new avenues in cancer treatment.

## Supporting information

Arrive Checklist

Z stack video

Z stack video

Uncropped gels and blots

## List of abbreviations

*ALDH1*: Aldehyde dehydrogenase 1
*ABCC1*: ATP binding cassette subfamily C member 1
*ABCB1*: ATP binding cassette subfamily C member 1
*BC*: Breast Cancer
*BCSCs*: Breast Cancer Stem Cells
*BCRP1*: Breast Cancer Resistance Protein
*CSCs*: Cancer Stem Cells
*EC*: Ehrlich Carcinoma
*EMT*: Epithelial to Mesenchymal Transition
*HE*: Haematoxylin Eosin
*Lin*: Lineage
*MDR*: Multidrug resistance
*MMP*: Matrix Metalloproteinase
*NT*: No treatment
*PSGL1*: P-Selectin Glycoprotein Ligand 1
*RP*: Resting Platelet
*TEP*: Tumor Educated Platelet
*TNBC*: Triple negative breast cancer
*TME*: Tumor microenvironment
*TCIPA*: Tumor cell induced platelet aggregation
*VEGF*: Vascular Endothelial Growth Factor
*VEGFR1*: Vascular Endothelial Growth Factor Receptor 1
*VEGFR2*: Vascular Endothelial Growth Factor Receptor 2
*WNT*: Wingless Related Integration Site

## Declarations

### Ethics approval and consent to participate

Human breast tumor and blood samples, were collected from Chittaranjan National Cancer Institute (CNCI), Kolkata, India following written consent from patients and Institutional Ethical Committee (Approval no: CNCI-IEC-SB-2020-30).

1. Title of the approved project: Elucidating the role of tumor educated platelets in promoting Epithelial to Mesenchymal transition and angiogenesis in breast cancer: Modulation by 2DG/NLGP
2. Name of the institutional approval committee or unit: Institutional Ethics Committee, Chittaranjan National Cancer Institute, Kolkata, India
3. Approval number: CNCI-IEC-SB-2020-30
4. Date of approval: December 4^th^, 2020

The human cell lines MCF-7 (RRID: CVCL_0031) and MDA-MB-231(RRID: CVCL_0062) were procured from National Centre for Cell Science, NCCS, Pune, India. Cell line preparation was permitted by the concerned Ethical Committee along with the human donors, through informed consents.

### Ethics approval for animal experiments

All experiments were performed after approval from Institutional Animal Care and Ethics Committee (Approval No: IAEC-1774/SBn-4/2021/9), registered under the Committee for the Purpose of Control and Supervision of Experiments on Animals (CPCSEA), Government. of India.

1. Title of the approved project: Elucidating the role of tumor educated platelets in promoting Epithelial to Mesenchymal transition and angiogenesis in breast cancer: Modulation by 2DG/NLGP
2. Name of the institutional approval committee or unit: Institutional Animal Care and Ethics Committee, registered under the Committee for the Purpose of Control and Supervision of Experiments on Animals (CPCSEA), Government. of India.
3. Approval number: IAEC-1774/SBn-4/2021/9
4. Date of approval: March 12^th^, 2021

### Consent for publication

All authors have given their consent for publication

### Availability of data and materials

No datasets were generated or analysed during the current study.

### Author’s contribution

**A. G, S. Banerjee, A. B** designed the experiments, **A. G** performed all the experiments, **A.G, S. Banerjee** analysed data, **A.G** prepared figures, **A. G** wrote and **S. Banerjee, A. Bose, R.N.B**. reviewed the manuscript, **J.S** assisted with cell culture, **S.B, P. Roy Choudhury** assisted with *in-vivo* experiments, **M.C, S.D, A.S, J.D, N.G, T.D, I. Guha** supported in preliminary experiments, **N.A, I.G, S.H** provided pre-operative patient blood and post-operative human tumor samples, **S. Banerjee, A. Bose, R.N.B** supervised the whole work.

## Acknowledgements

We would like to acknowledge the support of the Director, Chittaranjan National Cancer Institute, Kolkata, India for providing institutional facilities. Dr Abhijit Rakshit, Head, Animal Facility, CNCI and all lab members for their technical help for this work. We are especially thankful to Dr. Swapna Chaudhury, ICMR, Emeritus scientist, CNCI, Kolkata for her generous gift of antibodies. We are also thankful to Ms. Shalini Das and Mr. Rupankar Ghosh, Technical Support operator, BD Biosciences for acquisition of flow-cytometric data. We are much obliged to Dr. Sumit Kumar Hira, Assistant Professor, University of Burdwan, Dr. Priyajit Chatterjee, University Science Instrument Centre, University of Burdwan for their expertise in obtaining scanning electron microscope photographs. We also thank Mr. Soham Bose, Department of Biochemistry, University of Calcutta, for his support in confocal microscopy-based imaging.

**‘The authors declare that they have not used AI-generated work in this manuscript’.**

## References

1. Yersal O, Barutca S. Biological subtypes of breast cancer: Prognostic and therapeutic implications. World journal of clinical oncology. 2014 Aug 8;5(3):412.

2. Yin L, Duan JJ, Bian XW, Yu SC. Triple-negative breast cancer molecular subtyping and treatment progress. Breast Cancer Research. 2020 Dec; 22:1–3.

3. Guha A, Goswami KK, Sultana J, Ganguly N, Choudhury PR, Chakravarti M, Bhuniya A, Sarkar A, Bera S, Dhar S, Das J. Cancer stem cell–immune cell crosstalk in breast tumor microenvironment: a determinant of therapeutic facet. Frontiers in Immunology. 2023 Nov 27; 14:1245421.

4. Rahman M, Mohammed S. Breast cancer metastasis and the lymphatic system. Oncology letters. 2015 Sep 1;10(3):1233–9.

5. Ordaz-Ramos A, Tellez-Jimenez O, Vazquez-Santillan K. Signaling pathways governing the maintenance of breast cancer stem cells and their therapeutic implications. Frontiers in Cell and Developmental Biology. 2023;11.

6. Zhang X, Powell K, Li L. Breast cancer stem cells: biomarkers, identification and isolation methods, regulating mechanisms, cellular origin, and beyond. Cancers. 2020 Dec 14;12(12):3765.

7. Crabtree JS, Miele L. Breast cancer stem cells. Biomedicines. 2018 Jul 17;6(3):77.

8. Velasco-Velázquez MA, Popov VM, Lisanti MP, Pestell RG. The role of breast cancer stem cells in metastasis and therapeutic implications. The American journal of pathology. 2011 Jul 1;179(1):2–11.

9. Budczies J, von Winterfeld M, Klauschen F, Bockmayr M, Lennerz JK, Denkert C, Wolf T, Warth A, Dietel M, Anagnostopoulos I, Weichert W. The landscape of metastatic progression patterns across major human cancers. Oncotarget. 2015 Jan;6(1):570.

10. Ha NH, Faraji F, Hunter KW. Mechanisms of metastasis. Cancer Targeted Drug Delivery: An Elusive Dream. 2013 Jul 10:435–58.

11. Bian X, Yin S, Yang S, Jiang X, Wang J, Zhang M, Zhang L. Roles of platelets in tumor invasion and metastasis: a review. Heliyon. 2022 Dec 1;8(12).

12. Liao K, Zhang X, Liu J, Teng F, He Y, Cheng J, Yang Q, Zhang W, Xie Y, Guo D, Cao G. The role of platelets in the regulation of tumor growth and metastasis: the mechanisms and targeted therapy. MedComm. 2023 Oct;4(5): e350.

13. Lucotti S, Muschel RJ. Platelets and metastasis: new implications of an old interplay. Frontiers in oncology. 2020 Sep 18; 10:1350.

14. Assinger A. Platelets and infection–an emerging role of platelets in viral infection. Frontiers in immunology. 2014 Dec 18; 5:124104.

15. Zhou L, Zhang Z, Tian Y, Li Z, Liu Z, Zhu S. The critical role of platelet in cancer progression and metastasis. European Journal of Medical Research. 2023 Sep 28;28(1):385.

16. Najafi S, Asemani Y, Majidpoor J, Mahmoudi R, Aghaei-Zarch SM, Mortezaee K. Tumor-educated platelets. Clinica Chimica Acta. 2023 Dec 4:117690.

17. Schlesinger M. Role of platelets and platelet receptors in cancer metastasis. Journal of hematology & oncology. 2018 Oct 11;11(1):125.

18. In ‘t Veld SG, Wurdinger T. Tumor-educated platelets. Blood, The Journal of the American Society of Hematology. 2019 May 30;133(22):2359–64.

19. Antunes-Ferreira M, D’Ambrosi S, Arkani M, Post E, In ‘t Veld SG, Ramaker J, Zwaan K, Kucukguzel ED, Wedekind LE, Griffioen AW, Oude Egbrink M. Tumor-educated platelet blood tests for Non-Small Cell Lung Cancer detection and management. Scientific reports. 2023 Jun 8;13(1):9359.

20. Puricelli C, Boggio E, Gigliotti CL, Stoppa I, Sutti S, Giordano M, Dianzani U, Rolla R. Platelets, Protean Cells with All-Around Functions and Multifaceted Pharmacological Applications. International Journal of Molecular Sciences. 2023 Feb 26;24(5):4565.

21. Wang X, Zhao S, Wang Z, Gao T. Platelets involved tumor cell EMT during circulation: communications and interventions. Cell Communication and Signaling. 2022 Jun 3;20(1):82.

22. Zuo XX, Yang Y, Zhang Y, Zhang ZG, Wang XF, Shi YG. Platelets promote breast cancer cell MCF-7 metastasis by direct interaction: surface integrin α2β1-contacting-mediated activation of Wnt-β-catenin pathway. Cell Communication and Signaling. 2019 Dec;17:1–5.

23. Labelle M, Begum S, Hynes RO. Direct signaling between platelets and cancer cells induces an epithelial-mesenchymal-like transition and promotes metastasis. Cancer cell. 2011 Nov 15;20(5):576–90.

24. Ren Z, Liang S, Yang J, Han X, Shan L, Wang B, Mu T, Zhang Y, Yang X, Xiong S, Wang G. Coexpression of CXCR4 and MMP9 predicts lung metastasis and poor prognosis in resected osteosarcoma. Tumor Biology. 2016 Apr;37:5089–96.

25. Li Z, Chen G, Ding L, Wang Y, Zhu C, Wang K, Li J, Sun M, Oupicky D. Increased survival by pulmonary treatment of established lung metastases with dual STAT3/CXCR4 inhibition by siRNA nanoemulsions. Molecular Therapy. 2019 Dec 4;27(12):2100–10.

26. Yeeravalli R, Das A. Molecular mediators of breast cancer metastasis. Hematology/Oncology and Stem Cell Therapy. 2021 Dec 1;14(4):275–89.

27. Owyong M, Chou J, van den Bijgaart RJ, Kong N, Efe G, Maynard C, Talmi-Frank D, Solomonov I, Koopman C, Hadler-Olsen E, Headley M. MMP9 modulates the metastatic cascade and immune landscape for breast cancer anti-metastatic therapy. Life science alliance. 2019 Dec 1;2(6).

28. Gay LJ, Felding-Habermann B. Contribution of platelets to tumour metastasis. Nature Reviews Cancer. 2011 Feb;11(2):123–34.

29. Bhuniya A, Sarkar A, Guha A, Choudhury PR, Bera S, Sultana J, Chakravarti M, Dhar S, Das J, Guha I, Ganguly N. Tumor activated platelets induce vascular mimicry in mesenchymal stem cells and aid metastasis. Cytokine. 2022 Oct 1; 158:155998.

30. Varkey J, Nicolaides T. Tumor-educated platelets: A review of current and potential applications in solid tumors. Cureus. 2021 Nov 1;13(11).

31. Ding S, Dong X, Song X. Tumor educated platelet: the novel BioSource for cancer detection. Cancer Cell International. 2023 Dec;23(1):1–4.

32. Lin Y, Huang S, Qi Y, Xie L, Jiang J, Li H, Chen Z. PSGL-1 is a novel tumor microenvironment prognostic biomarker with cervical high-grade squamous lesions and more. Frontiers in Oncology. 2023 Mar 8;13:1052201.

33. Dimitroff CJ, Descheny L, Trujillo N, Kim R, Nguyen V, Huang W, Pienta KJ, Kutok JL, Rubin MA. Identification of leukocyte E-selectin ligands, P-selectin glycoprotein ligand-1 and E-selectin ligand-1, on human metastatic prostate tumor cells. Cancer research. 2005 Jul 1;65(13):5750–60.

34. Heidemann F, Schildt A, Schmid K, Bruns OT, Riecken K, Jung C, Ittrich H, Wicklein D, Reimer R, Fehse B, Heeren J. Selectins mediate small cell lung cancer systemic metastasis. PloS one. 2014 Apr 3;9(4):e92327.

35. Radziwon-Balicka A, Medina C, O’driscoll L, Treumann A, Bazou D, Inkielewicz-Stepniak I, Radomski A, Jow H, Radomski MW. Platelets increase survival of adenocarcinoma cells challenged with anticancer drugs: mechanisms and implications for chemoresistance. British journal of pharmacology. 2012 Oct;167(4):787–804.

36. Huong PT, Nguyen LT, Nguyen XB, Lee SK, Bach DH. The role of platelets in the tumor-microenvironment and the drug resistance of cancer cells. Cancers. 2019 Feb 19;11(2):240.

37. Pradella D, Naro C, Sette C, Ghigna C. EMT and stemness: flexible processes tuned by alternative splicing in development and cancer progression. Molecular cancer. 2017 Dec;16:1–9.

38. Roy S, Sunkara RR, Parmar MY, Shaikh S, Waghmare SK. EMT imparts cancer stemness and plasticity: new perspectives and therapeutic potential. Frontiers in Bioscience-Landmark. 2020 Oct 1;26(2):238–65.

39. Gawlik-Rzemieniewska N, Bednarek I. The role of NANOG transcriptional factor in the development of malignant phenotype of cancer cells. Cancer biology & therapy. 2016 Jan 2;17(1):1–0.

40. Khales SA, Mozaffari-Jovin S, Geerts D, Abbaszadegan MR. TWIST1 activates cancer stem cell marker genes to promote epithelial-mesenchymal transition and tumorigenesis in esophageal squamous cell carcinoma. BMC cancer. 2022 Dec 6;22(1):1272.

41. Chakravarti M, Dhar S, Bera S, Sinha A, Roy K, Sarkar A, Dasgupta S, Bhuniya A, Saha A, Das J, Banerjee S. Terminally exhausted CD8+ T cells resistant to PD-1 blockade promote generation and maintenance of aggressive cancer stem cells. Cancer Research. 2023 Jun 2;83(11):1815–33.

42. Douyère M, Chastagner P, Boura C. Neuropilin-1: a key protein to consider in the progression of pediatric brain tumors. Frontiers in Oncology. 2021 Jul 1;11:665634.

43. He K, Gan WJ. Wnt/β-catenin signaling pathway in the development and progression of colorectal cancer. Cancer Management and Research. 2023 Dec 31:435–48.

44. Hwang I, Kim J, Jeong S. β-Catenin and peroxisome proliferator-activated receptor-δ coordinate dynamic chromatin loops for the transcription of vascular endothelial growth factor A gene in colon cancer cells. Journal of Biological Chemistry. 2012 Nov 30;287(49):41364–73.

45. Kwon IK, Schoenlein PV, Delk J, Liu K, Thangaraju M, Dulin NO, Ganapathy V, Berger FG, Browning DD. Expression of cyclic guanosine monophosphate-dependent protein kinase in metastatic colon carcinoma cells blocks tumor angiogenesis. Cancer. 2008 Apr 1;112(7):1462–70.

46. Easwaran V, Lee SH, Inge L, Guo L, Goldbeck C, Garrett E, Wiesmann M, Garcia PD, Fuller JH, Chan V, Randazzo F. β-Catenin regulates vascular endothelial growth factor expression in colon cancer. Cancer research. 2003 Jun 15;63(12):3145–53.

47. Lichtenberger LM, Vijayan KV. Are platelets the primary target of aspirin’s remarkable anticancer activity? Cancer research. 2019 Aug 1;79(15):3820–3.

48. . Pulcinelli FM, Pignatelli P, Celestini A, Riondino S, Gazzaniga PP, Violi F. Inhibition of platelet aggregation by aspirin progressively decreases in long-term treated patients. Journal of the American College of Cardiology. 2004 Mar 17;43(6):979–84.

49. Undas A, Brummel-Ziedins KE, Mann KG. Antithrombotic properties of aspirin and resistance to aspirin: beyond strictly antiplatelet actions. Blood. 2007 Mar 15;109(6):2285–92.

50. Ornelas A, Zacharias-Millward N, Menter DG, Davis JS, Lichtenberger L, Hawke D, Hawk E, Vilar E, Bhattacharya P, Millward S. Beyond COX-1: the effects of aspirin on platelet biology and potential mechanisms of chemoprevention. Cancer and Metastasis Reviews. 2017 Jun; 36:289–303.

51. DeRogatis JM, Viramontes KM, Neubert EN, Tinoco R. PSGL-1 immune checkpoint inhibition for CD4+ T cell cancer immunotherapy. Frontiers in Immunology. 2021 Feb 23; 12:636238.

