## Supplementary Figures S1-S8 for "Tumor-Educated-Platelets interact with Breast Cancer-Stem-Cells via P-selectin- PSGL1 and ensure stemness and metastasis through WNT-β-Catenin-VEGF-VEGFR2 intra-cellular signaling: Therapeutic modulation by aspirin"

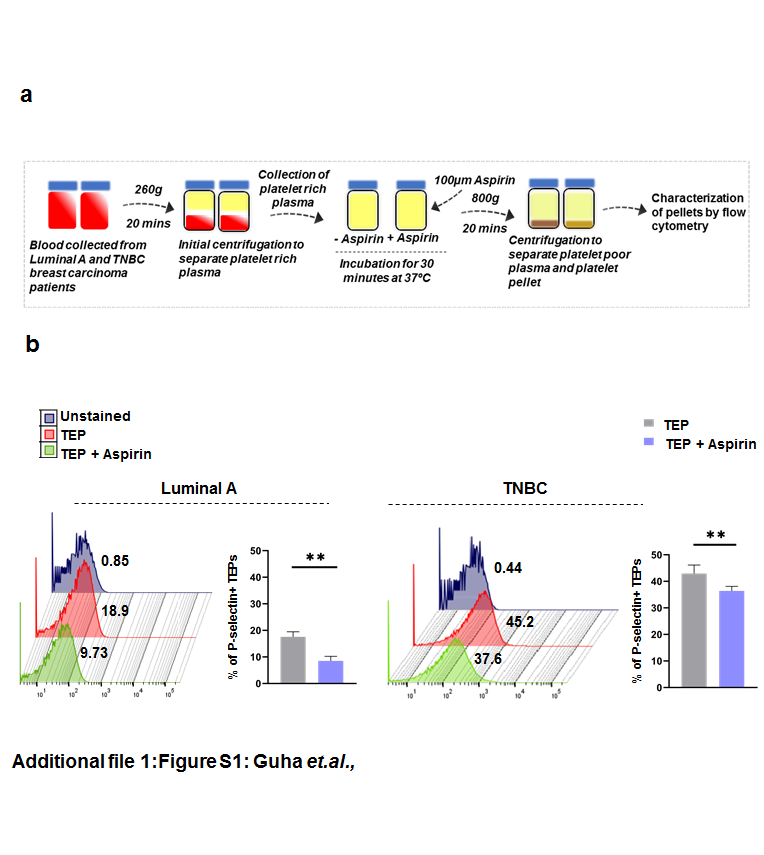

**Supplementary Figure S1: Guha *et.al.,***

**Supplementary Figure S2: Guha *et.al.,***

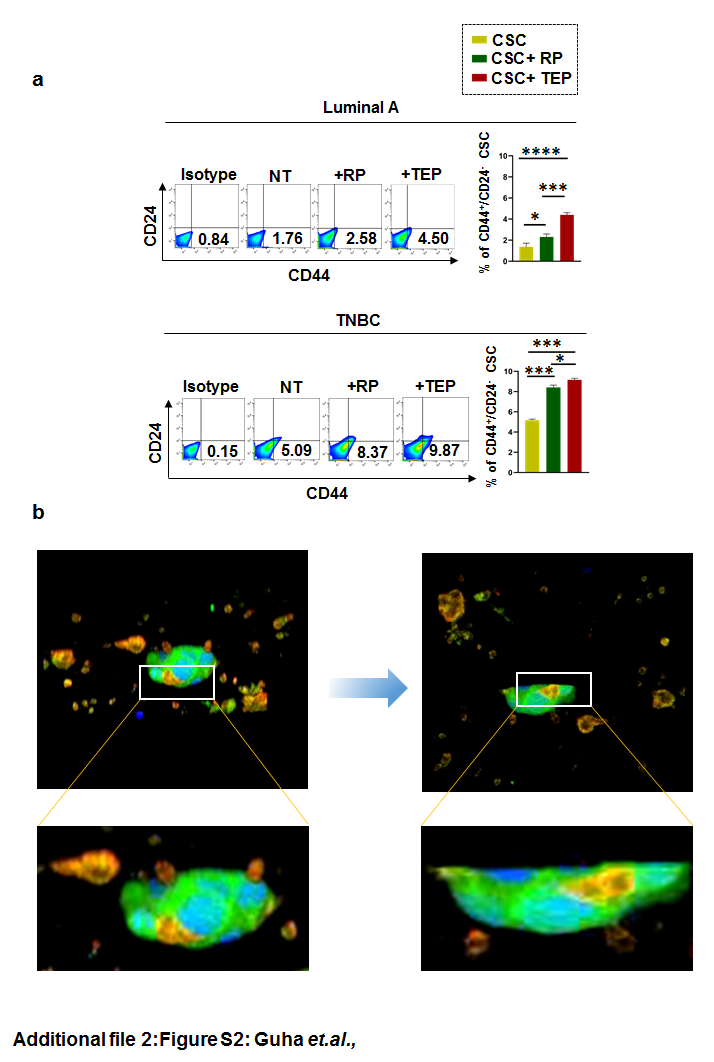

**Supplementary Figure S2: Guha *et.al.,***

**Supplementary Figure S3: Guha *et.al.,***

**Supplementary Figure S3: Guha *et.al.,***

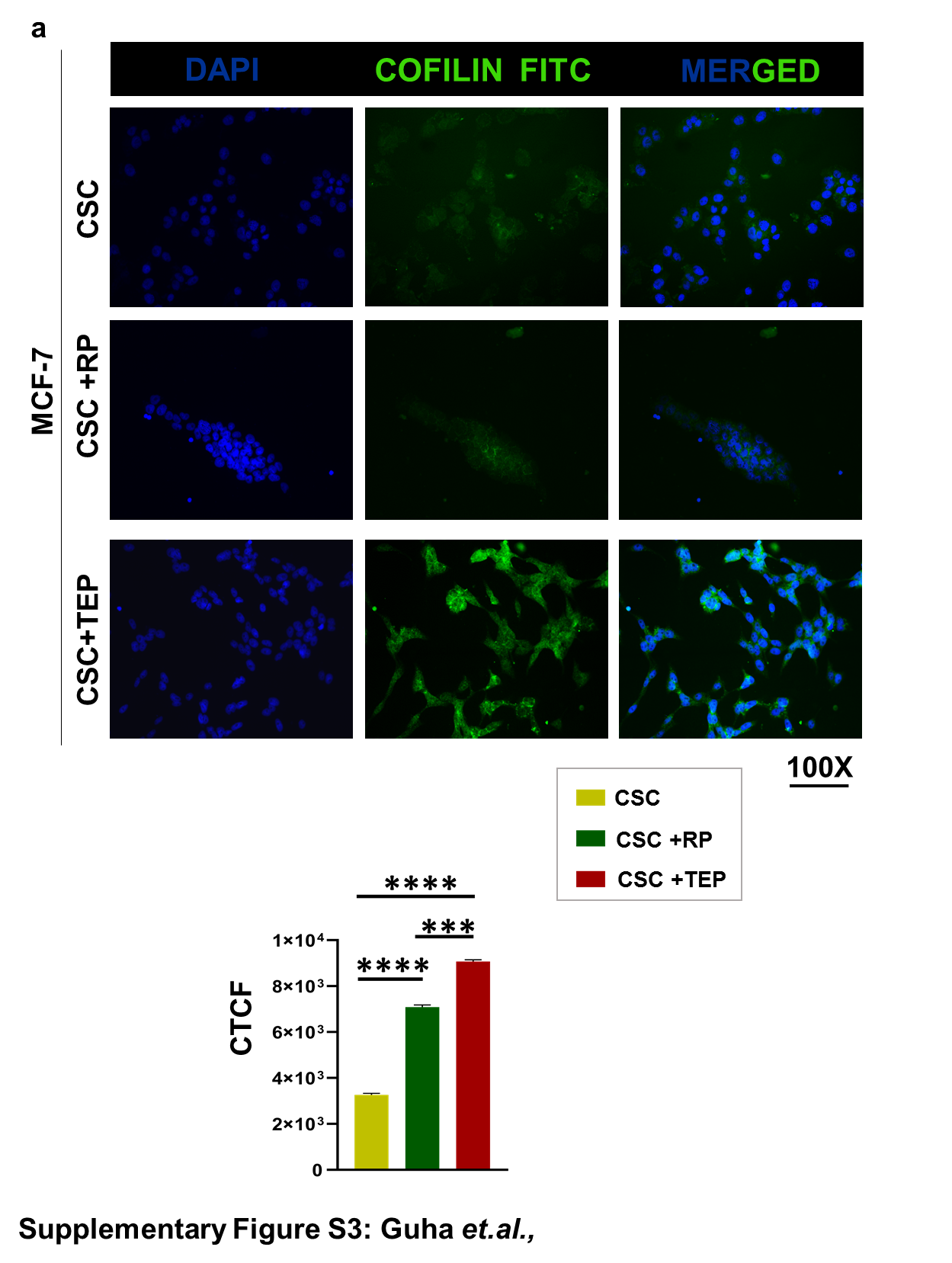

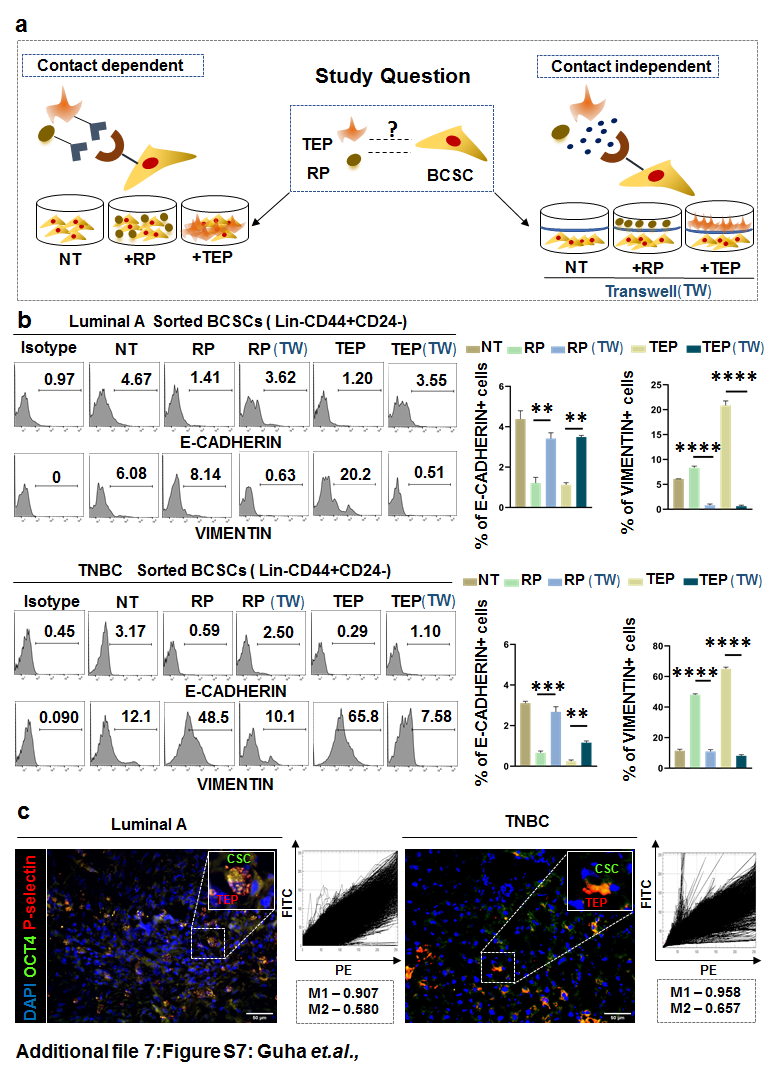

**Supplementary Figure S4: Guha *et.al.,***

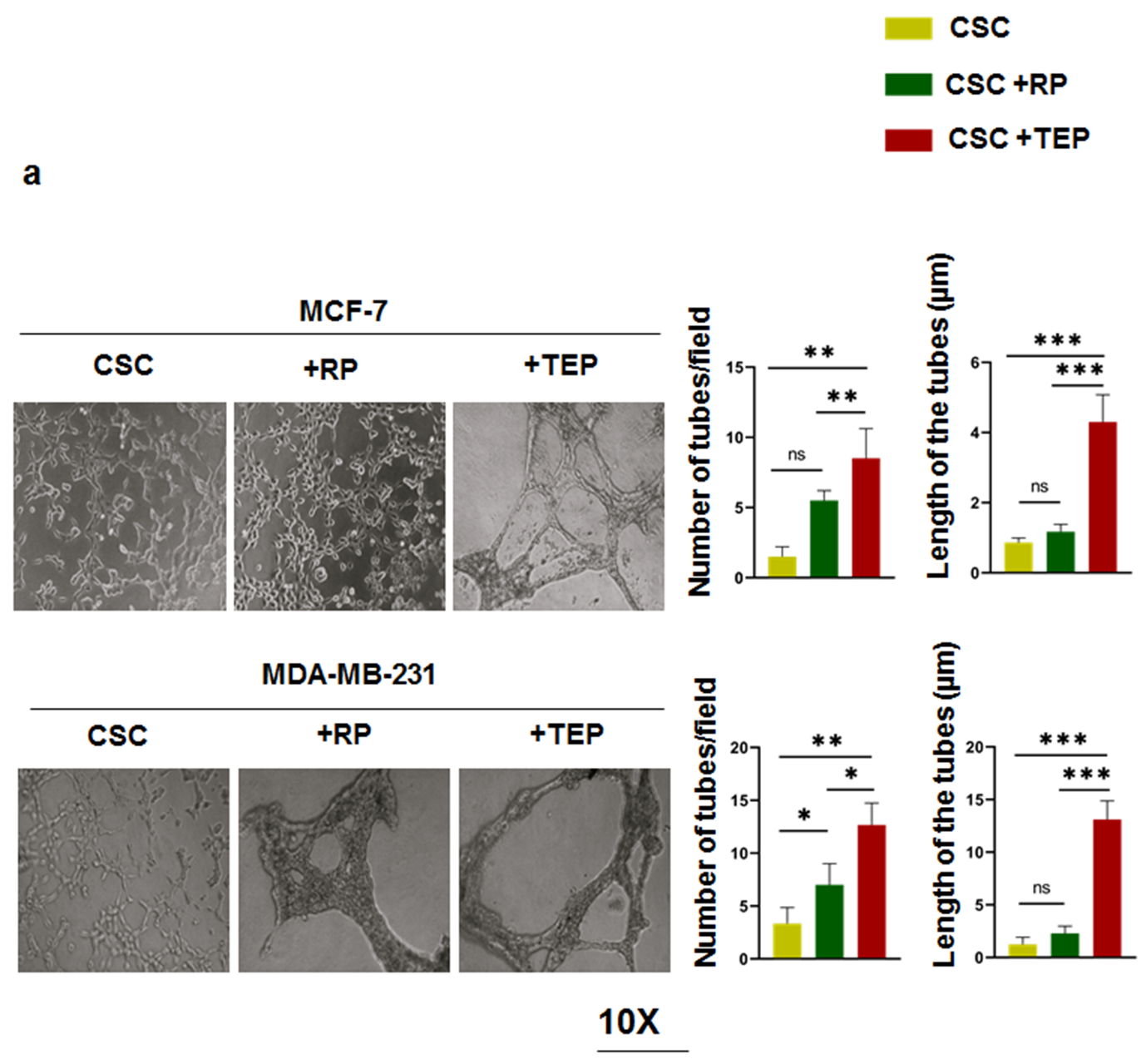

**Supplementary Figure S5: Guha *et.al.,***

**Supplementary Figure S5: Guha *et.al.,***

**Supplementary Figure S6: Guha *et.al.,***

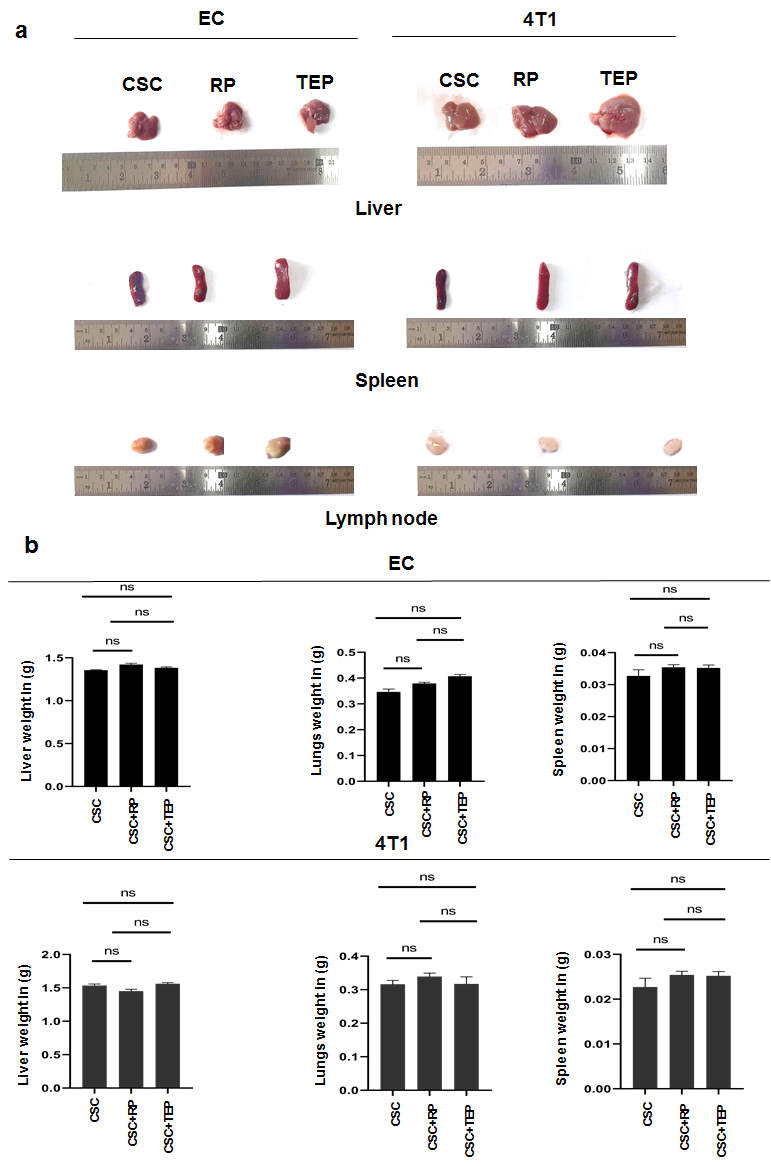

**Supplementary Figure S6: Guha *et.al.,***

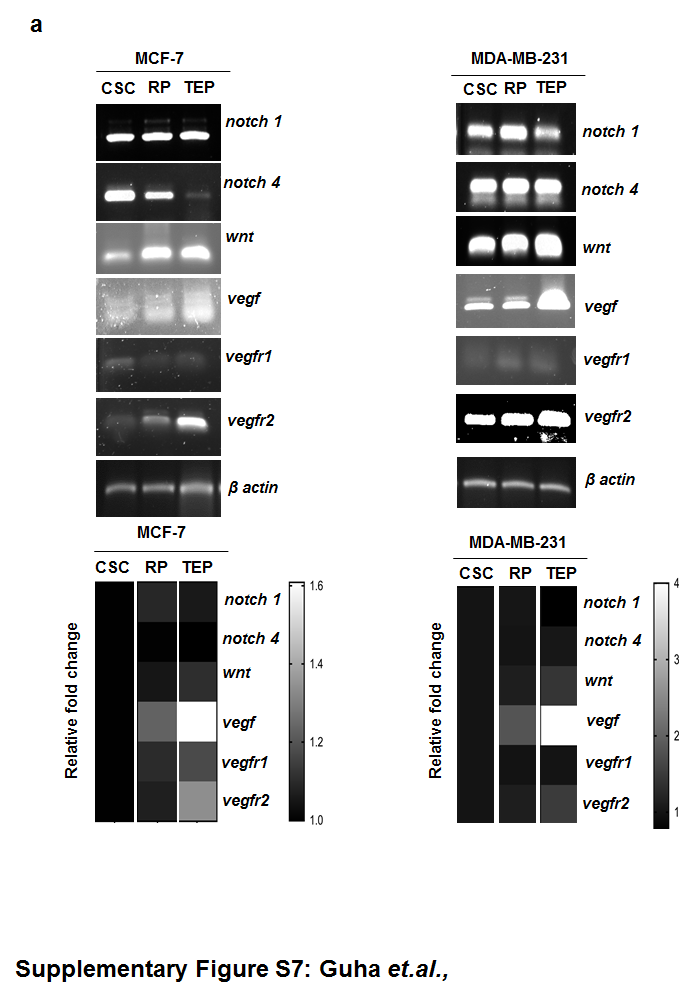

**Supplementary Figure S7: Guha *et.al.,***

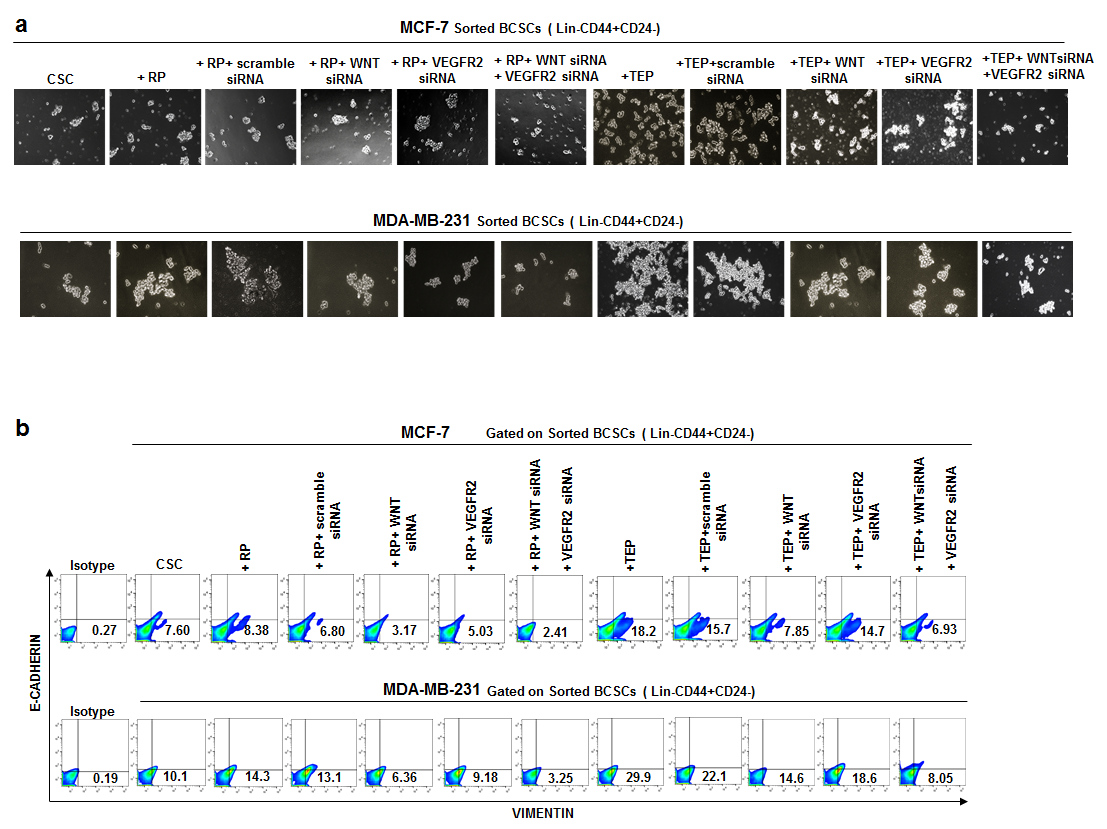

**Supplementary Figure S8: Guha *et.al.,***

**Supplementary Figure S1: a.** Schematic depiction of isolation and treating TEPs with aspirin. **b.** Flow cytometric histogram plots depicting reduction in frequency of P- selectin^+^ TEPs upon treatment with aspirin in both luminal A and TNBC. In bar-graph, (mean±SD) of TEP% is presented with statistical significance inferred from unpaired non-parametric t-test, followed by two-tailed p value was performed (n=20). *Inset* blue, red and green bar denoting unstained, TEP, TEP + aspirin population*p<0.05, **p<0.01, ***p<0.001, ****p<0.0001, ns: not significant are indicated.

**Supplementary Figure S2: a.** Pseudo-colour flow cytometric plots portraying changes in CD44^+^24^-^ BCSC frequency in CSC, CSC+RP, CSC+TEP cohorts of luminal A and TNBC. Bar diagram illustrating % of CD44^+^24^-^ BCSCs keeping (mean±SD) across all experimental groups. Statistical significance is drawn from one-way ANOVA followed by Tukey’s multiple comparison test (n=6).

**Supplementary Figure S3: a.** Representative immunofluorescence images at 100X magnification of CSC, CSC+RP, CSC+TEP of MCF-7 stained with cofilin FITC. Quantified corrected total cell fluorescence intensity (CTCF) is displayed in bar-graph, (mean±SD) with statistical significance inferred from one-way ANOVA followed by Tukey’s multiple comparison test (n=3). *Inset*: yellow, green and red column denoting CSC, CSC+RP, CSC+TEP. *p<0.05, **p<0.01, ***p<0.001, ****p<0.0001, ns: not significant are indicated.

**Supplementary Figure S4: a.** Schematic illustration of probable contact-dependent and contact-independent interactions between BCSCs +/- TEP/RP in luminal A and TNBC subtypes. **b.** Representative flow-cytometric histograms of E-CADHERIN and VIMENTIN expression of BCSCs isolated from co-culture setups of contact-dependent and contact-independent arrangements of all the groups across luminal A and TNBC sub-types. Bar graphs (mean±SD) depicting changes in expression E-CADHERIN and VIMENTIN in contact-dependent and independent setups of all the groups in both the cohorts is presented. Statistical significance is drawn from one-way ANOVA followed by Tukey’s multiple comparison test (n=3). *p<0.05, **p<0.01, ***p<0.001, ****p<0.0001, ns: not significant are indicated. **c.** Representative immunofluorescence images of breast tumor sections of luminal A and TNBC subtypes at 40X magnification depicting proximity of BCSCs (stained with OCT-FITC) and TEPs (stained with P-selectin PE). In *inset* enlarged image of their close adjacency is presented. Cytofluoromicrographs and Mander’s co-localization coefficient (M1- overlap of PE over FITC and M2- overlap of FITC over PE) of both the sub-types is presented.

**Supplementary Figure S5: a.** Illustrative micrographs at 10X magnification of interconnected tube formation on matrigel across the groups of both MCF-7 and MDA-MB-231. In bar-graph, (mean±SD) of number of tubes/field and tube length in µm is given. Statistical significance is drawn from one-way ANOVA followed by Tukey’s multiple comparison test (n=3). *Inset*: yellow, green and red column denoting CSC, CSC+RP, CSC+TEP. *p<0.05, **p<0.01, ***p<0.001, ****p<0.0001, ns: not significant are indicated.

**Supplementary Figure S6: a.** Illustrative photographs of unaffected murine liver and spleen and lymph node of experimental metastasis model of CSC, CSC+RP, CSC+TEP of both EC and 4T1. **b.** Weight in grams of liver and spleen and lymph node of EC (upper panel) and 4T1 (lower panel) murine models is given. In bar-graph, (mean±SD) of organ weight is provided with statistical significance inferred from one-way ANOVA followed by Tukey’s multiple comparison test (n=6).

**Supplementary Figure S7: a.** mRNA expression of *notch1, notch4, wnt, vegf, vegfr1, vegfr2* in CSC, CSC+RP, CSC+TEP of MCF-7 and MDA-MB-231 by RT-PCR is represented keeping β actin as control (n=3). Relative fold change in expression levels is provided by heatmaps. Darker colour denotes lower expression while lighter colour signifies higher expression respectively. Uncropped full-length gel is provided in Additional file 5.

**Supplementary Figure S8: a.** Representative micrographs of mammospheres at 10X magnification across all experimental groups in presence and absence of WNT siRNA and VEGFR2 siRNA in MCF-7 and MDA-MB-231. **b.** Pseudo-colour flow cytometric plots depicting changes in CD44^+^24^-^VIMENTIN^+^E-CADHERIN^-^metastatic BCSC frequency across all experimental cohorts of MCF-7 and MDA-MB-231 in presence and absence of WNT siRNA and VEGFR2 siRNA.
