## Supplementary Tables for "Tumor-Educated-Platelets interact with Breast Cancer-Stem-Cells via P-selectin- PSGL1 and ensure stemness and metastasis through WNT-β-Catenin-VEGF-VEGFR2 intra-cellular signaling: Therapeutic modulation by aspirin"

**Supplementary Table T1-**

**Reagent Antibody Details:**

| Sl. No | Antibody/Antibody-cocktail kit | Make | Catalogue |
| --- | --- | --- | --- |
| 1. | Purified CD41 Antibody | Genetex | GTX113758 |
| 2. | Ultra-leaf purified anti-human-CD62P | Biolegend | 304947 |
| 3. | APC-mouse anti human CD62P | BD Biosciences | 561920 |
| 4. | Purified PSGL1 Antibody | Santa Cruz | SC-10172 |
| 5. | Purified anti ALDH1L1 Antibody | Biolegend | 856802 |
| 6. | Lineage Cell Depletion Kit, human | Miltenyi Biotec | 130-092-211 |
| 7. | CD44 MicroBeads, human | Miltenyi Biotec | 130-095-194 |
| 8. | CD24 MicroBead Kit, human | Miltenyi Biotec | 130-095-951 |
| 9. | FITC anti-mouse/human CD44 Antibody | Biolegend | 103005 |
| 10. | PE anti-mouse/human CD24 Antibody | Biolegend | 101807 |
| 11. | Purified Nanog Antibody | Santa Cruz | SC-374103 |
| 12. | Purified SOX2 Antibody | R & D Systems | MAB2018 |
| 13. | Purified OCT4 Antibody | R & D Systems | MAB1759 |
| 14. | Purified Anti-mouse E-cadherin | Novus | NBP2-19051 |
| 15. | Purified Anti-mouse Vimentin | R & D Systems | MAB2105 |
| 16. | Purified Twist Antibody | Genetex | GTX127310 |
| 17. | Purified Cofilin Antibody | Cell Signaling Technology | 5175 |
| 18. | Purified WNT Antibody | Santa Cruz | SC-6266 |
| 19. | Purified β-catenin Antibody | Elabscience | ENT0672 |
| 20. | Purified Frizzled Antibody | Santa Cruz | SC-7429 |
| 21. | Purified VEGF Antibody | Santa Cruz | SC-507 |
| 22. | Purified Flt1 (VEGFR1) | Santa Cruz | SC-316 |
| 23. | Purified Anti-Flk-1/KDR/VEGFR2 Antibody | Santa Cruz | SC-6251 |
| 24. | HRP- Anti-Mouse IgG Antibody | Sigma | A5278 |
| 25. | HRP-Anti-Rat IgG Antibody | Abcam | ab6734 |
| 26. | HRP-Anti-Rabbit IgG Antibody | Sigma | A0545 |
| 27. | HRP-Anti-Goat IgG Antibody | Sigma | A5420 |
| 28. | FITC Anti-Rabbit IgG Antibody | Santa Cruz | SC-2012 |
| 29. | PE Goat Anti-Mouse IgG Antibody | BD Pharmingen | 550589 |
| 30. | Donkey Anti-Goat IgG PE |  | SC-3743 |
| 31. | PerCP-Cy 5.5 Anti-Mouse Antibody | BD Biosciences | 560668 |
| 32. | APC/Cy7 Anti-Mouse Antibody | Abcam | AB130785 |
| 33. | FITC Anti-Mouse IgG Antibody | Sigma | F5897 |
| 34. | TRITC - Anti-Rat Antibody | Abcam | ab6841 |
| 35. | Goat Anti-Rabbit FITC Antibody | Santa Cruz | SC 2012 |
| 36. | Goat Anti-Rat FITC Antibody | Abcam | AB6840 |

**Supplementary Table T2-**

**Primer Details:**

| SL. No | Primer Name | Forward primer sequence 5’- 3’ | Reverse primer sequence |
| --- | --- | --- | --- |
| 1. | Human *β-actin* | 5’-AGCGAGCATCCCCCAAAGTT-3’ | 5’-GGGCACGAAGGCTCATCATT-3’ |
| 2. | Human *bcrp1* | 5’-TCAGGAGGCCTTGGGATACT-3’ | 5’-AGTTCCACGGCTGAAACACT-3’ |
| 3. | Human *abcb1* | 5’-ATTTTCAATGTTTCGCTATT-3’ | 5’-TTCATGAAGAACCCTGTATC-3’ |
| 4. | Human *abcc1* | 5’-TCGTGTGGGTGCCTTGTTT-3’ | 5’-AACAGCAGCACGGTGTAGAA-3’ |
| 5. | Human *snail* | 5’-CTTCGTCCTTCTCCTCTACT-3’ | 5’-ATTCCTTGTTGCAGTATTTG-3’ |
| 6. | Human *slug* | 5’-TACAGTCCAAGCTTTCAGAC-3’ | 5’-GCTCACATATTCCTTGTCAC-3’ |
| 7. | Human *ccr7* | 5’-ATACTAGCCACAGTGCTCTC-3’ | 5’-GAAGACTATGAAGACCACGA-3’ |
| 8. | Human *cxcr3* | 5’-ATGGAGTTGAGGAAGTACG-3’ | 5’-CACTCTCGTTTTCTCCATAG-3’ |
| 9. | Human *cxcr4* | 5’-ACCAACAGTCAGAGGCCAAG-3’ | 5’-ACACAACCACCCACAAGTCA-3’ |
| 10. | Human *cxcr5* | 5’-CCATGCTCTACACTTTCGCC-3’ | 5’-AGAACGTGGTGAGAGAGGTG-3’ |
| 11. | Human *mmp7* | 5’-GAGTGCCAGATGTTGCAGAA-3’ | 5’-AAATGCAGGGGGATCTCTTT-3’ |
| 12. | Human *mmp9* | 5’-TTGACAGCGACAAGAAGTGG-3’ | 5’-GCCATTCACGTCGTCCTTAT-3’ |
| 13. | Human *mmp11* | 5’-TAGGTGCCTGCATCTGTCTG-3’ | 5’-TGGCTTTGGAGGATAGCAGT-3’ |
| 14. | Human *wnt* | 5’-AGAGAGGGTAGAAGACGTTG-3’ | 5’-GAGGAACACTGACCTAGTCC-3’ |
| 15. | Human *notch1* | 5’-GAACTGTGAGGAAAATATCG-3’ | 5’-GACACACACGCAGTTGTAG-3’ |
| 16. | Human *notch4* | 5’-AGAAAGACTCCACCTTTCAC-3’ | 5’-GTCTCACACTCATCCACATC-3’ |
| 17. | Human *vegf* | 5’-GGCCTCCGAAACCATGAACT-3’ | 5’-GCTGCGCTGATAGACATCCA-3’ |
| 18. | Human *vegfr1* | 5’-CTGGGCAGCAGACAAATCCT-3’ | 5’-CACAACCAAGGTGCTAGCCA-3’ |
| 19. | Human *vegfr2* | 5’-GTACACCTGTGCAGCATCCA-3’ | 5’-AATCGTCAGTACATGCCCCG-3’ |

### Supplementary Table T3-

**Human patient sample details:**

| Sl.No | AGE | Hormone Receptor Status | TNM Stage |
| --- | --- | --- | --- |
| 1. | 56 | TNBC | T4N1MX |
| 2. | 50 | TNBC | T3N1MX |
| 3. | 52 | TNBC | T2N2MX |
| 4. | 58 | TNBC | T2N0MX |
| 5. | 56 | TNBC | T4N1MX |
| 6. | 50 | TNBC | T3N1MX |
| 7. | 52 | TNBC | T2N2MX |
| 8. | 58 | TNBC | T2N0M0 |
| 9. | 54 | TNBC | T2N1M0 |
| 10. | 50 | TNBC | T2N1M0 |
| 11. | 58 | TNBC | T2N0M0 |
| 12. | 52 | TNBC | T2N0M0 |
| 13. | 54 | TNBC | T2N1M0 |
| 14. | 53 | TNBC | T3NIM0 |
| 15. | 60 | TNBC | T2N1MX |
| 16. | 58 | TNBC | T4N1M0 |
| 17. | 51 | TNBC | T4N1MX |
| 18. | 55 | TNBC | T2N1MX |
| 19. | 58 | TNBC | T2N0M0 |
| 20. | 48 | TNBC | T3N1MX |
| 21. | 60 | Luminal A | T4N1MX |
| 22. | 62 | Luminal A | T2N1MX |
| 23. | 45 | Luminal A | T2N1MX |
| 24. | 54 | Luminal A | T3N1M0 |
| 25. | 55 | Luminal A | T4N1MX |
| 26. | 41 | Luminal A | T4N2M0 |
| 27. | 56 | Luminal A | T3N2M1 |
| 28. | 48 | Luminal A | T2N1MX |
| 29. | 67 | Luminal A | T2N1MX |
| 30. | 66 | Luminal A | T3N0MX |
| 31. | 59 | Luminal A | T4N1MX |
| 32. | 58 | Luminal A | T4N1M2 |
| 33. | 41 | Luminal A | T2N0MX |
| 34. | 52 | Luminal A | T4N1M1 |
| 35. | 48 | Luminal A | T2N1MX |
| 36. | 50 | Luminal A | T2N0M0 |
| 37. | 47 | Luminal A | T2N1MX |
| 38. | 51 | Luminal A | T2N0M0 |
| 39. | 51 | Luminal A | T2N1MX |
| 40. | 41 | Luminal A | T4N1M2 |
