## Supplementary material for "Tumor-Educated-Platelets interact with Breast Cancer-Stem-Cells via P-selectin- PSGL1 and ensure stemness and metastasis through WNT-β-Catenin-VEGF-VEGFR2 intra-cellular signaling: Therapeutic modulation by aspirin": Uncropped gels and blots

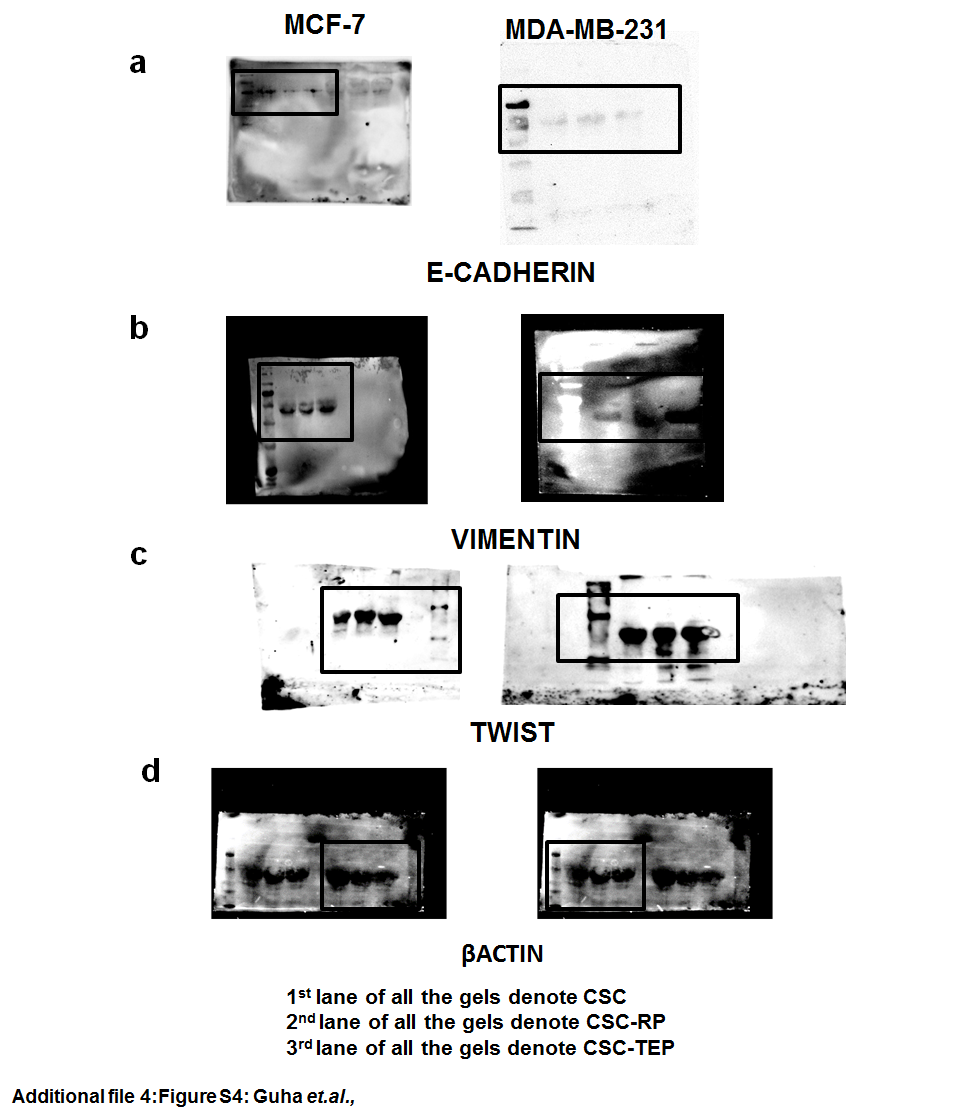

**Uncropped Raw blots with ladder for Figure 4b**

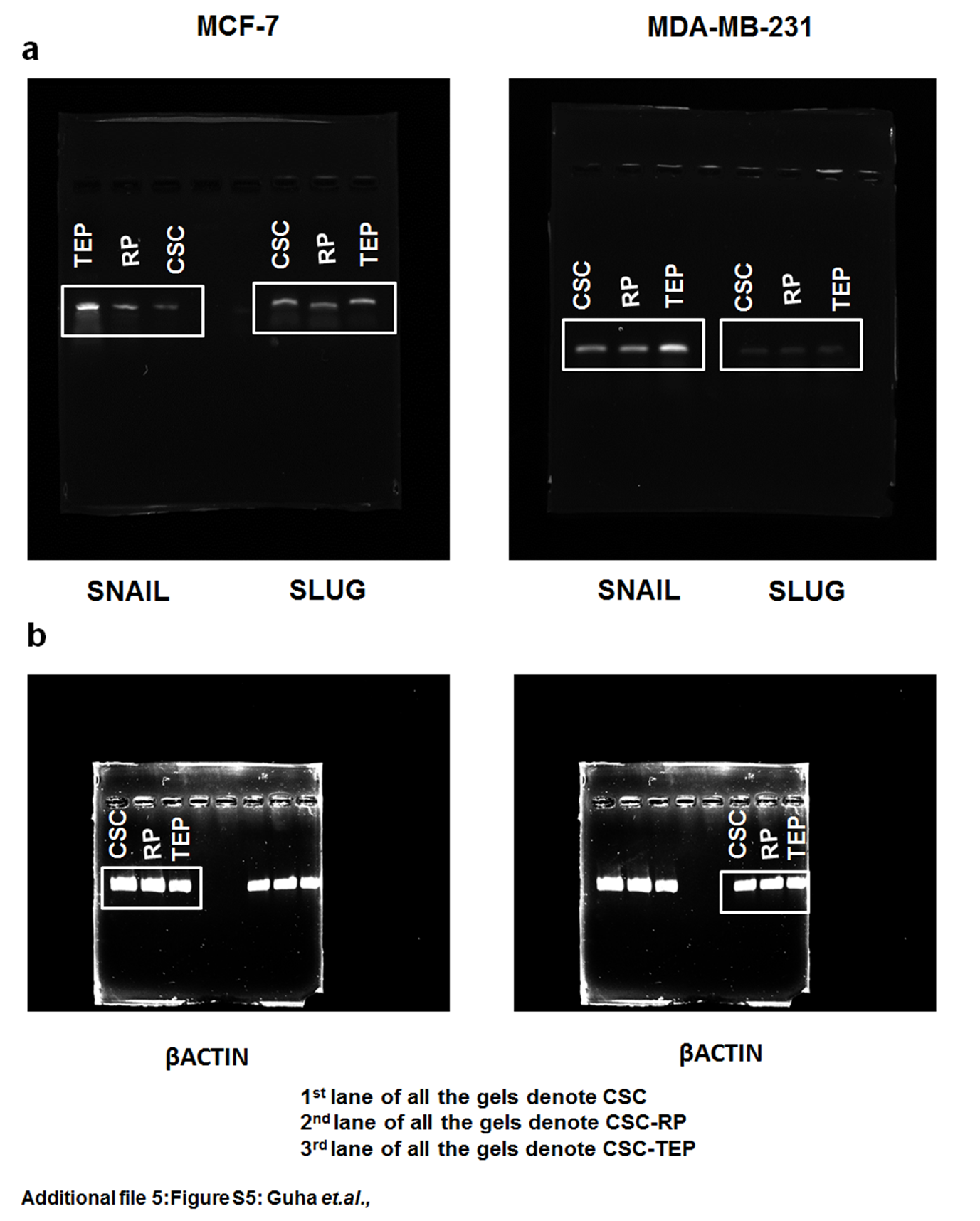

**Uncropped Raw gels for Figure 4c**

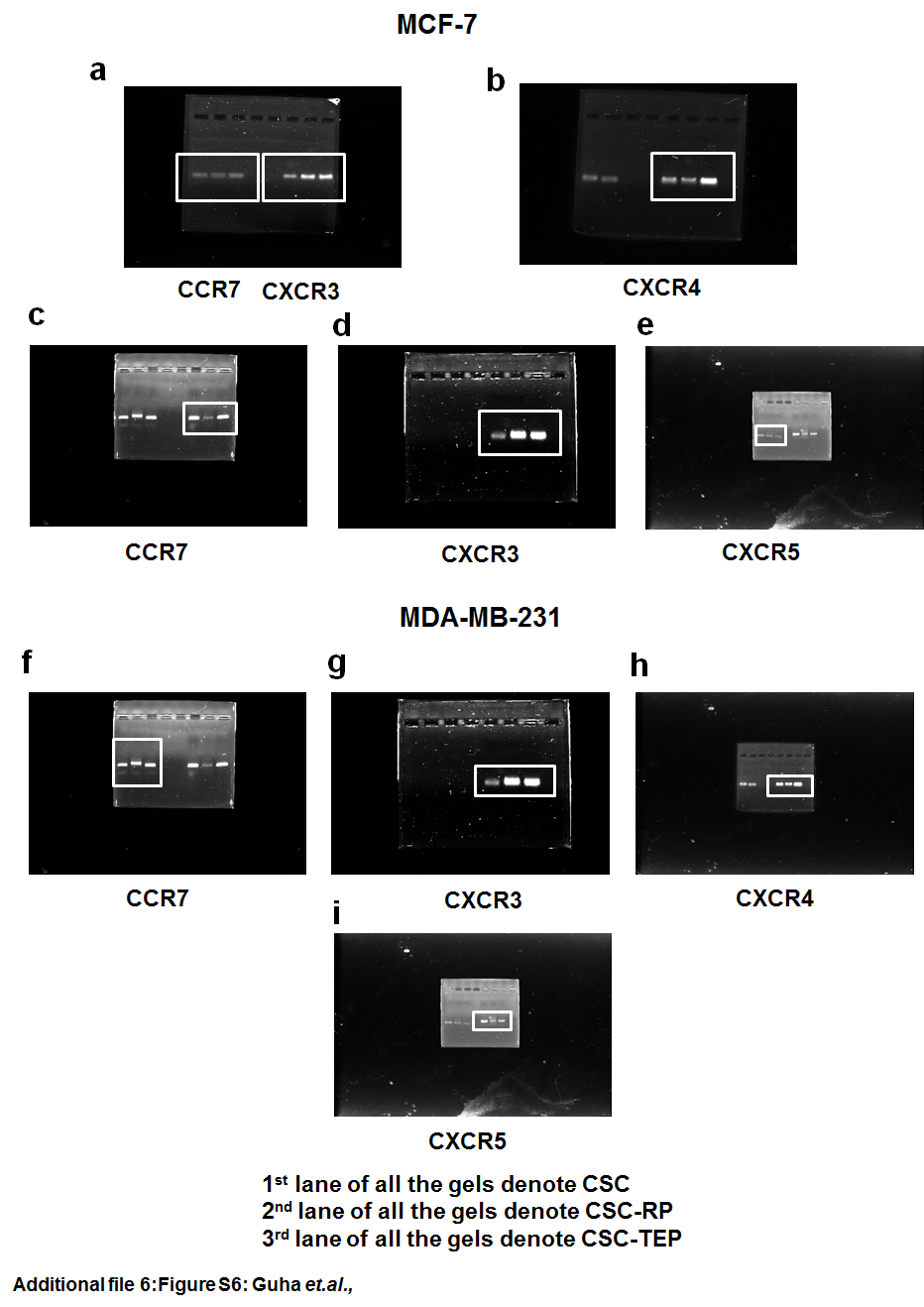

**Uncropped Raw gels for Figure 4d**

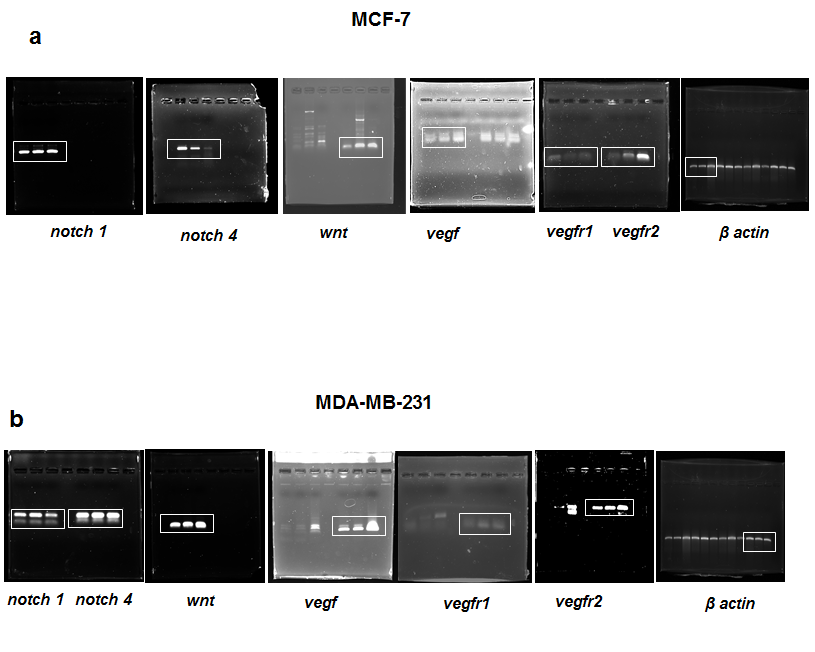

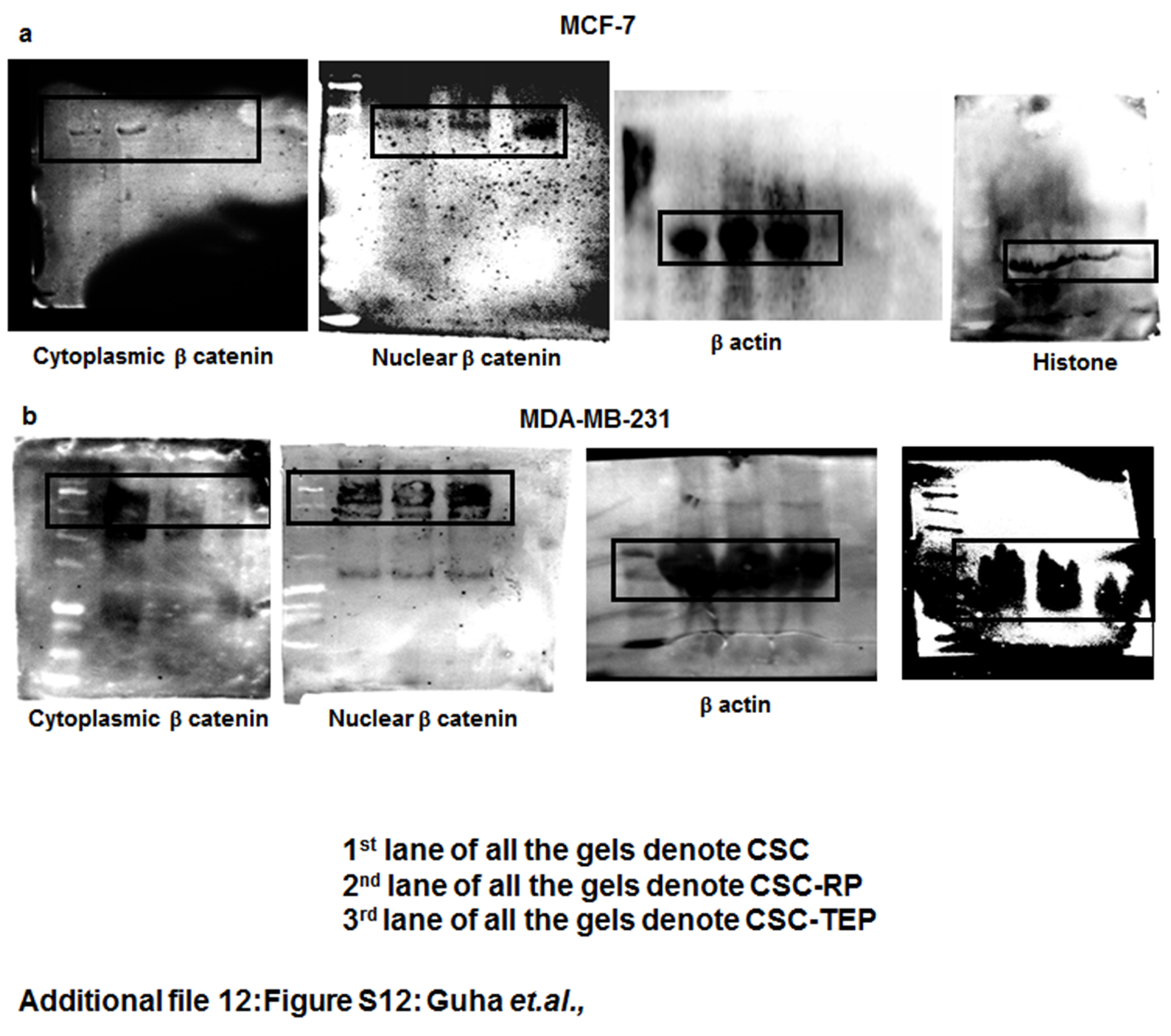

**Uncropped Raw gels for Supplementary Figure S7**

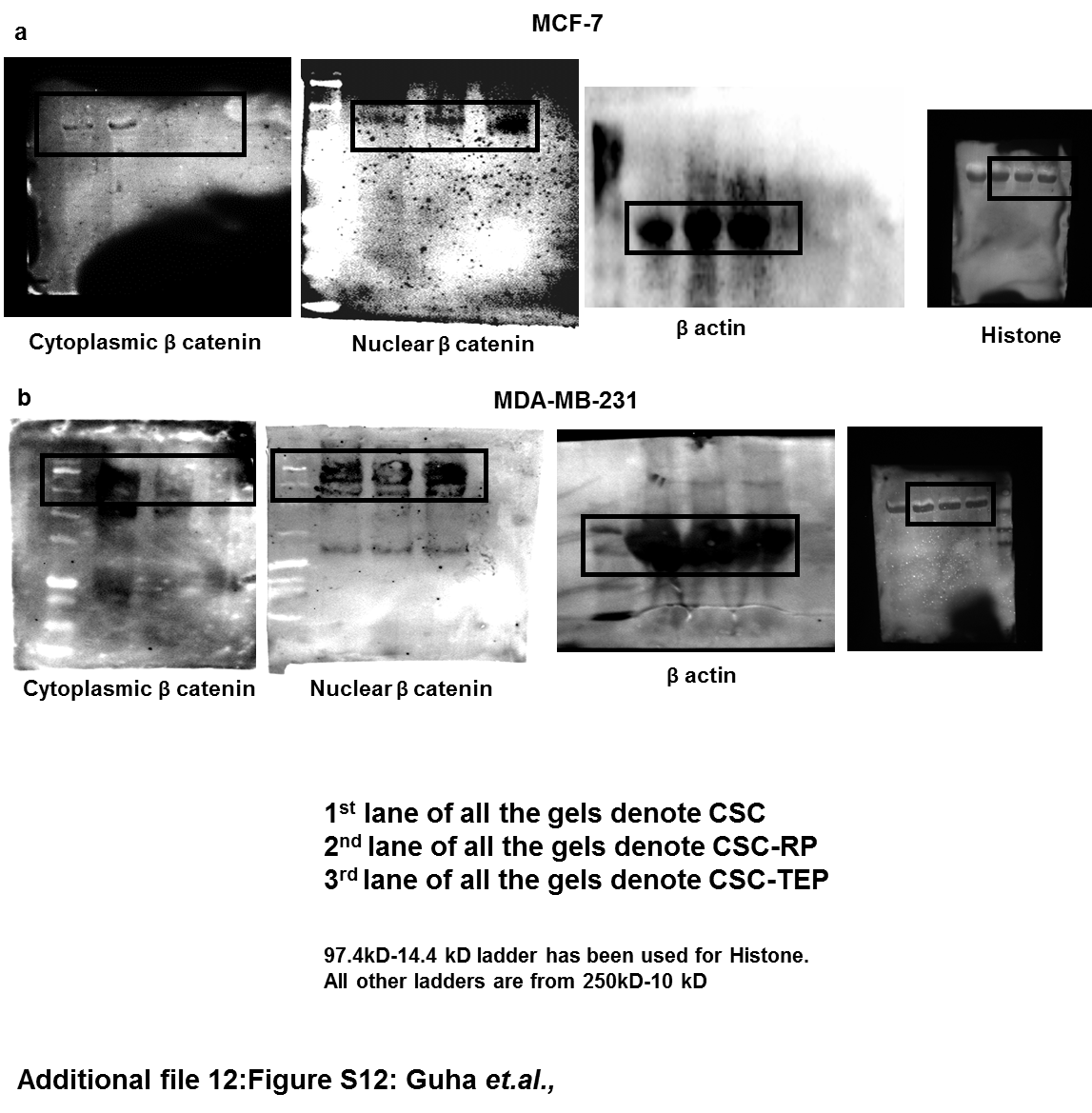

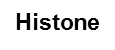

**Uncropped Raw blots with ladder for Figure 6c**
